# Identification of SPP1-positive macrophages by single-cell spatial analysis in human lung tissues with mycobacterial infection

**DOI:** 10.1101/2024.09.12.612778

**Authors:** Harutaka Katano, Akira Hebisawa, Yuko Sato, Yoshihiko Hoshino

## Abstract

Tuberculosis and non-tuberculous mycobacterial (NTM) diseases are infections caused by *Mycobacterium tuberculosis* and non-tuberculous mycobacteria such as the *Mycobacterium avium* complex, leading to the formation of granulomatous lesions with caseous necrosis in the lungs. Although granulomatous tissues are infiltrated by numerous inflammatory cells, including macrophages, lymphocytes, and neutrophils, the mechanisms underlying granuloma formation caused by mycobacteria remain unclear. In this study, we performed single-cell spatial analysis on lung tissue samples from patients with tuberculosis and NTM diseases to investigate the infiltrating cell populations. We analyzed seven lung lesions and identified individual cell types infiltrating the granulomatous tissue. Based on gene expression profiles, at least four macrophage subtypes were identified. Notably, SPP1-positive macrophages predominantly found infiltrating the granulomatous tissue. Langhans giant cells expressed SPP1, and numerous SPP1-positive macrophages without giant cell morphology were also observed around the granulomas. RNA-seq analysis revealed elevated SPP1 expression in mycobacterium-infected tissues. The SPP1-CD44 signaling pathway was active in SPP1-positive macrophages and their neighboring cells in mycobacterium-infected tissues. SPP1-positive macrophages were also observed around granulomas in other granulomatous diseases, such as granulomatosis with polyangiitis and sarcoidosis. These findings suggest that SPP1-CD44 signaling in SPP1-positive macrophages may play a role in the pathology of granulomatous diseases, including mycobacterial infections.

**Brief summary:** SPP1-CD44 signaling in SPP1-positive macrophages may play a role in granuloma formation in mycobacterial and other granulomatous diseases.

## INTRODUCTION

Tuberculosis and non-tuberculous mycobacterial (NTM) diseases, caused by *Mycobacterium tuberculosis* and various non-tuberculous mycobacteria—particularly the *Mycobacterium avium intracellulare* complex (MAC)—represent significant public health challenges(*1*). These infections predominantly affect the lungs, leading to the formation of granulomatous lesions, which are characterized by caseous necrosis(*2*). Despite extensive research efforts, the mechanisms underlying granuloma formation in mycobacterial infections remain largely enigmatic and poorly understood(*3–5*).

Granulomas are highly organized structures that form in response to persistent pathogens. These cellular conglomerates, composed of immune cells such as macrophages, lymphocytes, and neutrophils, aim to contain and neutralize the pathogen, serving as a barrier to prevent its spread(*2, 4, 6*). Immunohistochemical and special staining techniques have detected mycobacterial antigens within the caseous necrotic tissue of granulomas, supporting the hypothesis that granulomas play a crucial role in controlling the dissemination of mycobacteria(*3*). However, the precise formation and dynamic regulation of these structures, particularly in the context of mycobacterial infections, remain incompletely understood.

Various immune cells, including macrophages, B cells, T cells, NK cells, dendritic cells, and neutrophils, participate in granuloma formation.(*2, 7*). Among these, macrophages are pivotal, exhibiting remarkable plasticity that allows them to adapt to a wide range of environmental stimuli. During mycobacterial infections, macrophages can assume diverse functional states, contributing both to pathogen clearance and, paradoxically, to pathogen survival(*3, 4, 8, 9*). Macrophages, along with dendritic cells and alveolar epithelial cells, phagocytize mycobacteria during the initial stages of infection. The immune response to mycobacteria triggers several signaling pathways, including IFN-γ and TLR2 signaling, which promote the production of pro-inflammatory cytokines such as IL-12, IL-23, and TNF-α. These signals, particularly through TLR2, ultimately contribute to the formation of granulomas.(*8, 10–13*). To gain insight into the cellular responses to mycobacterial infections in the lung, it is critical to identify the specific cell types infiltrating granulomas and to characterize their dynamic interactions. To this end, we employed the cutting-edge Nanostring CosMx™ platform to perform single-cell spatial analysis on lung tissue samples from patients with MTB or NTM diseases. This advanced technology enabled us to map and identify individual cell types surrounding the granulomatous tissue(*14*). In addition, single-cell spatial analysis allowed for detailed gene expression profiling within individual cells, providing insights into the localization of cells and the identification of key signaling pathways involved in cell-cell communication within the tissue microenvironment. This innovative approach holds promise for unraveling the complex interplay of immune cells during MTB and NTM infections. In our study, we applied single-cell spatial analysis to explore the cellular landscape of lung tissue samples from patients with MTB or NTM infections, with a particular emphasis on macrophages and their critical role in granuloma formation.

## RESULTS

### Cell typing in lung tissues with mycobacterial infection

Seven lung tissue samples from patients with MTB (n = 3) and NTM (n = 4) infections were analyzed using the Nanostring CosMx™ platform (Fig. 1A and table S1). Each slide contained granulomas with inflammatory cell infiltrates, including foreign body macrophages (Langhans giant cells, figs. S1a and S1b). Immunohistochemistry and Ziehl-Neelsen staining demonstrated that mycobacteria bacilli were predominantly localized at the center of granulomas and absent from the surrounding inflammatory cell infiltrates in both NTM and MTB samples (fig. S2). The CosMx™ platform enables multiplex gene quantification for 960 genes in formalin-fixed paraffin-embedded (FFPE) slides with a documented sensitivity of 1-2 gene copies per cell and spatial transcript localization precision of less than 50 nm, corresponding to single cell resolution (*14*). CosMx™ analysis identified a total of 300,841 cells across the seven slides (table S3). After cell segmentation, transcription signals by in situ hybridization were detected in each cell on the slide (average: 131 transcripts per cell, Fig. 1B and table S3). All cells were clustered based on transcript expression profiles. Differentially expressed gene (DEG) analysis identified 520 cluster-specific marker genes with adjusted p-values. A total of 26 distinct clusters were identified (Fig. 1C and fig. S3A), which were visualized using uniform manifold approximation and projection (UMAP, Fig. 1D and fig. S3B). These DEGs (gene lists available in Supplementary Data: marker_heatmap_nb_clus_4.csv) were used to annotate subpopulations. The cell types within each cluster were determined by SingleR. Neutrophil and B-cell lineages were manually annotated based on gene expression data from the Human Gene Atlas (Gene Set Enrichment Analysis, GSEA, https://www.gsea-msigdb.org/gsea/index.jsp). All cells in each focus of view (fov) from the seven slides were annotated and labeled by colors in spatial images (Fig. 1E).

**Fig. 1.**
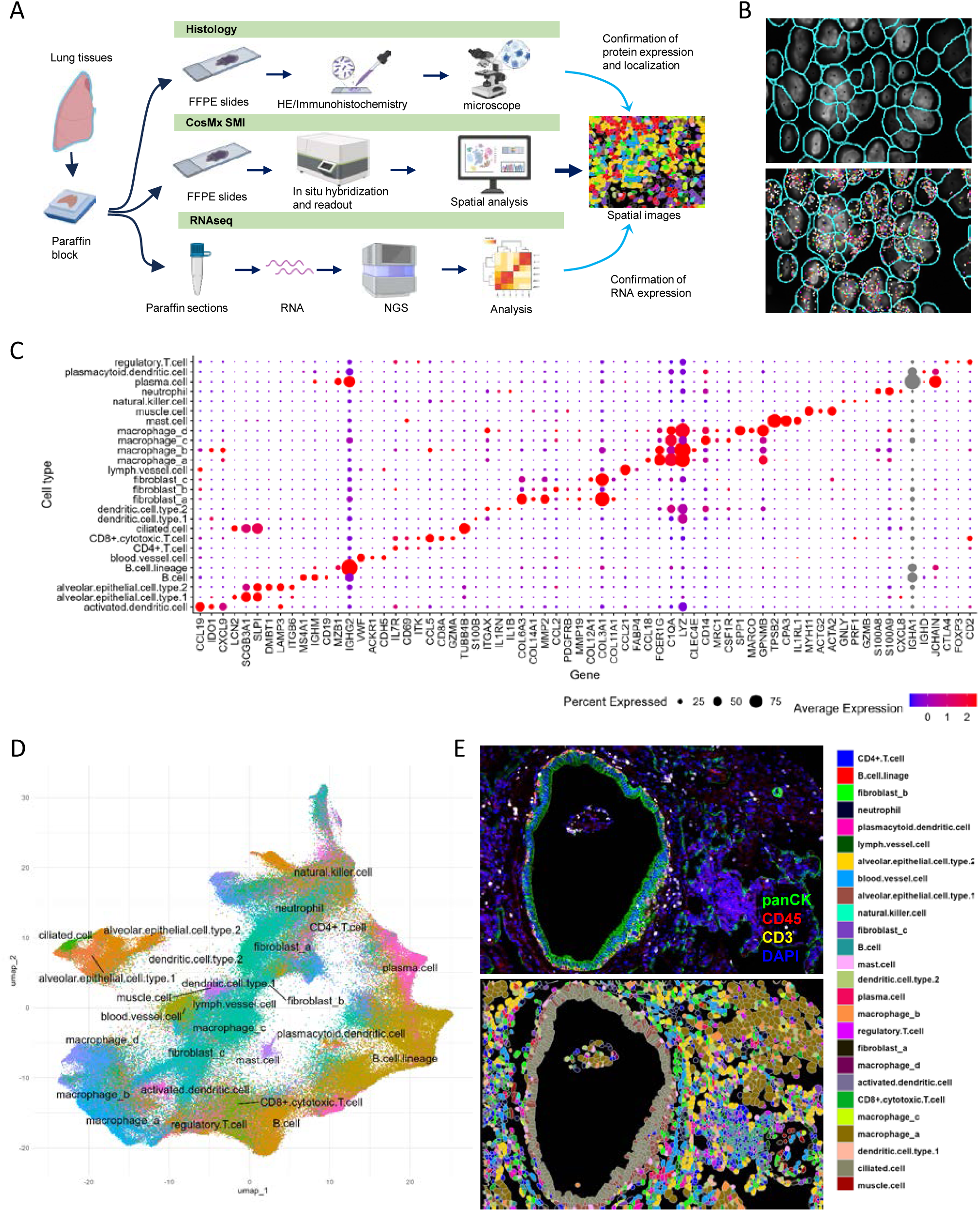
Single cell analysis of CosMx™ in mycobacterium-infected tissues. A. Overview of the study design for CosMx™ SMI, RNA-seq, and histological analysis. Lung samples, including MTB, NTM, granulomatous diseases and controls, were analyzed using RNA-seq (n = 37 samples) and CosMx™ SMI (n = 7 samples). This figure was created with BioRender.com. B. Representative image of CosMx™ SMI. All cells were segmented by segmentation markers and nuclei (upper panel). Transcripts (dots in various colors) were detected by in situ hybridization in each cell on the slides (lower panel). C. Dot plots showing expression of cell-typing markers in the indicated cell clusters. Dot size represents the percentage of cells expressing each gene in the cluster, while the average expression intensity of the markers is shown using a blue-to-red color gradient. D. UMAP plots of 300,841 cells from lung tissues with 4 NTM or 3 MTB patients. Each cluster is shown in a different color. E. Immunofluorescence image (upper panel) and spatial image obtained by CosMx™ SMI for MTB1 fov18 (lower panel). Cell annotations in the spatial image are listed in the right.

Through in silico analysis based on gene expression, 3 types of T cells were identified as CD4+, CD8+ cytotoxic, and regulatory T cells (fig. S4A). Three B-cell clusters were identified as B cells, plasma cells and B-cell linage cells. The cluster of B-cell lineage cluster exhibited gene expression patterns similar to plasma cells with J-chain expression being higher in plasma cells than B-cell lineage cluster (fig. S4B). Fibroblasts were categorized into three subtypes (a-c) based on varying collagen expression levels, including COL1A1 and COL3A1 (fig. S4C). Four macrophage subtypes (a-d) were identified, categorized based on the expression of specific markers such as CD14, CD163, CD68, and LYZ. Subtypes *a*, *b*, and *c* corresponding to M1 macrophages, M2 macrophage with low CD14 expression, and M2 macrophage with high CD14 expression, respectively (Figs. 2A and 2B). Macrophage subtype *d*, which expressed CCR7 and lacked CD163 expression (indicating an M1 phenotype), predominantly expressed SPP1 and MARCO (Figs. 2A-D). In violin, dot plots, and UMAP (Fig. 2), SPP1 expression appeared more specific to macrophage subtype d than MARCO and FN1, leading to its designation as SPP1-positive macrophages. The proportions of cell types varied across samples and FoVs (Fig. S5, Table S4). B cells (including B cells, B-cell lineage cells, and plasma cells) and fibroblasts (subtypes a–c) were the most abundant cell types, comprising 12.25% and 21.94% of the total cell population, respectively. Macrophages (subtypes a–d) and T cells (CD4+, CD8+, and Treg) accounted for 14.88% and 9.55% of the total cells, respectively. SPP1-positive macrophages (subtype d) represented 3.18% of all cells across the slides.

**Fig. 2.**
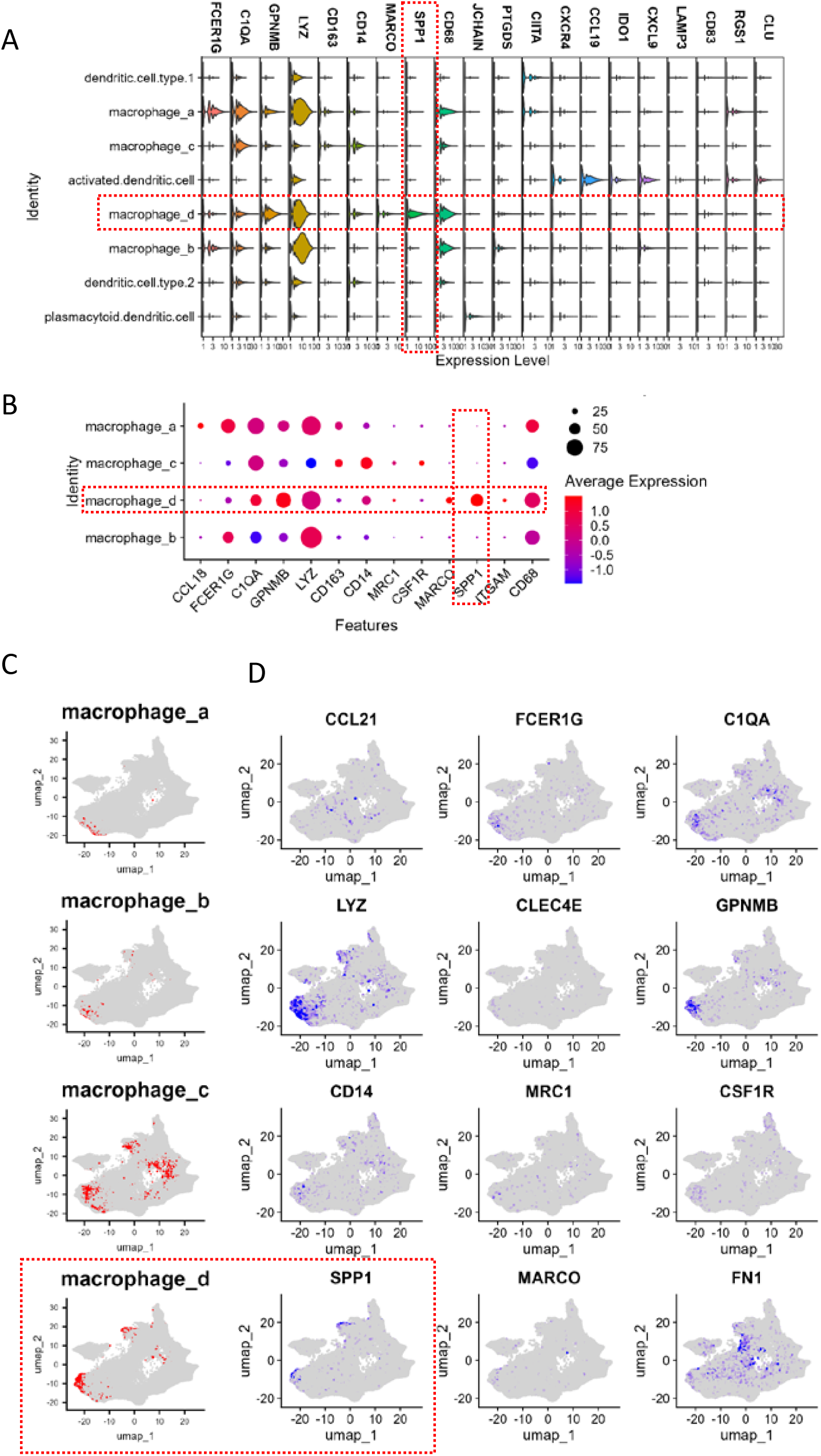
Subtyping of macrophages. A. Violin plot showing macrophage-associated gene expression in macrophages and dendritic cells. B. Dot plot illustrating macrophage-associated gene expression specially in macrophages. C. UMAP plot depicting four macrophage subtypes. D. UMAP plot displaying the expression of macrophage-associated genes.

### SPP1-positive macrophages infiltrate around MTB and NTM granulomas

CosMx™ spatial imaging revealed the cell types around granulomas. While numerous CD4 and CD8 T cells, B cells and macrophages infiltrated to interstitial regions and area around granulomas, macrophages were predominantly detected in the regions adjacent to granulomas (Fig. 3). Notably, SPP1-positive macrophages predominantly infiltrated the periphery of NTM-positive granulomatous tissue (Fig. 3 and figs. S6a and S6b). In MTB-infected tissues, SPP1-positive macrophages were also observed in the interstitial regions and alveolar spaces (Fig. 4 and fig. S6c). Immunohistochemistry using an anti-SPP1 antibody demonstrated that SPP1 expression was localized in the cytoplasm of Langhans giant cells (Figs. 3C and 4C), suggesting that these giant cells represent a form of SPP1-positive macrophages. Furthermore, CosMx™ spatial imaging and immunohistochemistry indicated that SPP1 was present in macrophages lacking giant cell morphology, located around mycobacterium-positive granulomas in close proximity to Langhans giant cells (Figs. 3A–3C). This observation suggests that SPP1-positive macrophages without giant cell morphology may share a similar origin with Langhans giant cells.

**Fig. 3.**
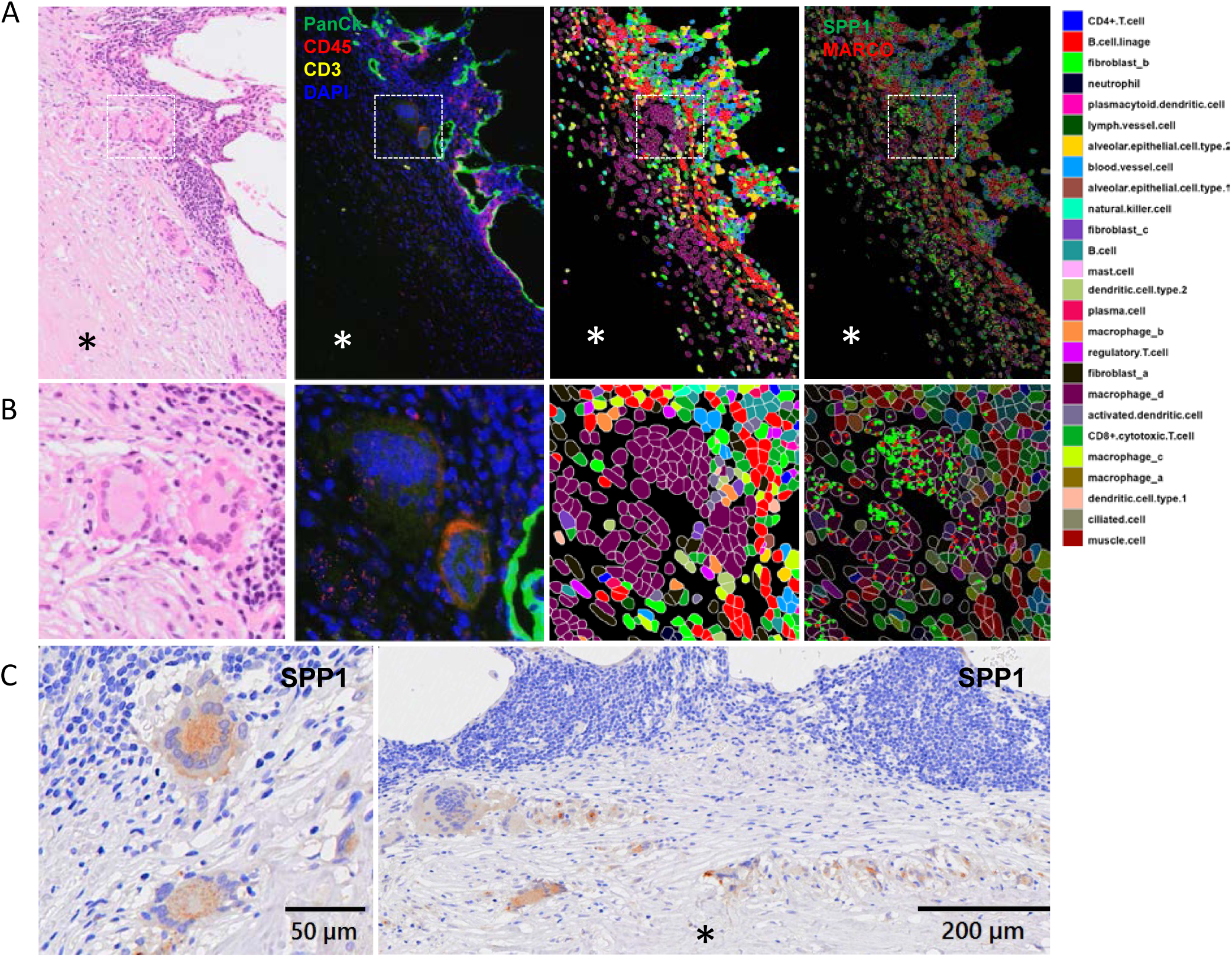
Spatial images of SPP1-positive macrophages in MAC-positive lung tissues. (Case NTM2) A. HE staining (left panel), immunofluorescence (second panel), and CosMx™ spatial images (right two panels) of macrophage_d. Colors representing different cell types are shown in the right panel. In CosMx™ images, SPP1 and MARCO signals are indicated by green and red dots, respectively. Asterisk indicates granulomatous region. B. Enlarged images of the boxed regions in A. C. Immunohistochemistry for SPP1 in the same tissue near the granuloma. Asterisk indicates granulomatous region.

**Fig. 4.**
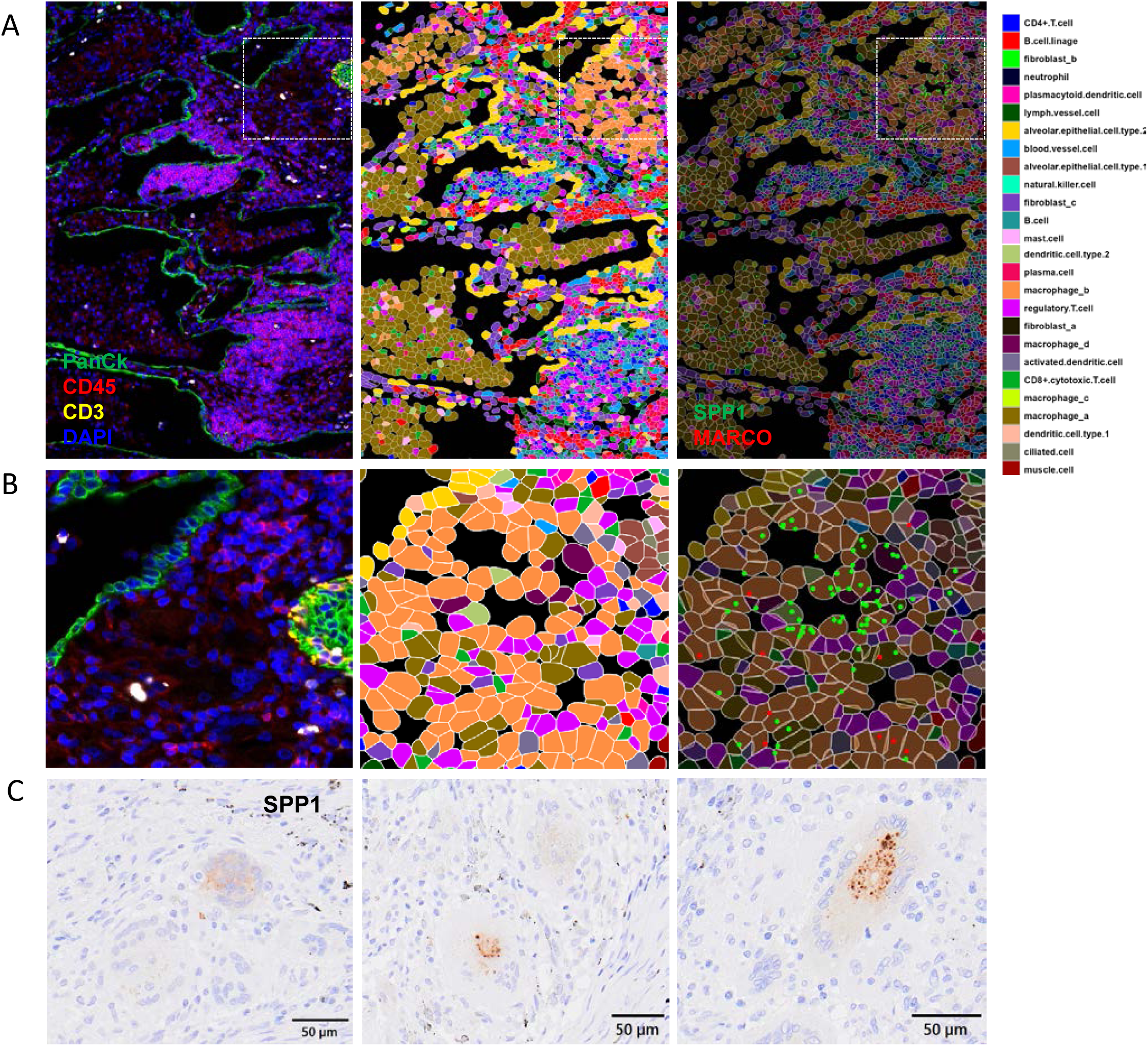
CosMx™ spatial images of SPP1 and MARCO expression in macrophage_d in MTB tissue. (Case MTB1) A. Macrophage_d in the interstitial region expressed SPP1 and MARCO. B. Enlarged images of the boxed regions in A. C. Immunohistochemistry for SPP1 in the same tissue near the granuloma.

Bulk RNA-seq analysis of RNA extracted from mycobacterium-infected tissues also revealed elevated expression of SPP1 in infected tissues compared to controls (figs. S7A-7C). Gene ontology analysis indicated that cytokine-mediated signaling pathways and extracellular matrix organization were upregulated in mycobacterium-infected samples (fig. S7D). These processes involved high expression of SPP1 and CD44, respectively.

### Ligand-receptor interaction of SPP1

Using spatial transcriptome data, the R package InSituCor swas employed for spatial correlation analysis to detect significant ligand-receptor interactions in the samples (fig. S8). This analysis identified SPP1, FN1, and MARCO as co-expression markers of SPP1-positive macrophages (Fig. 5A). Furthermore, InSituCor detected SPP1-CD44 and FN1-CD44 pairs as active ligand-receptor interactions in both NTM and MTB samples (Fig. 5B). Correlation network analysis using InSituCor identified numerous molecules involved in the SPP1-CD44 network (Fig. 5C). Spatial imaging of these samples demonstrated that CD44-positive cells were localized near SPP1-positive macrophages; however CD44-positive cells were distributed more broadly compared to SPP1-positive macrophages (fig. S9). Spatial analysis further revealed co-expression of CD44 and SPP1 in SPP1-positive macrophages (macrophage_d) within tissues infected with mycobacteria (Figs. 5D and 5E). Co-expression of SPP1 and CD44, as well as SPP1-CD44 ligand-receptor signaling, was frequently detected around granulomatous regions (fig. S10).

**Fig. 5.**
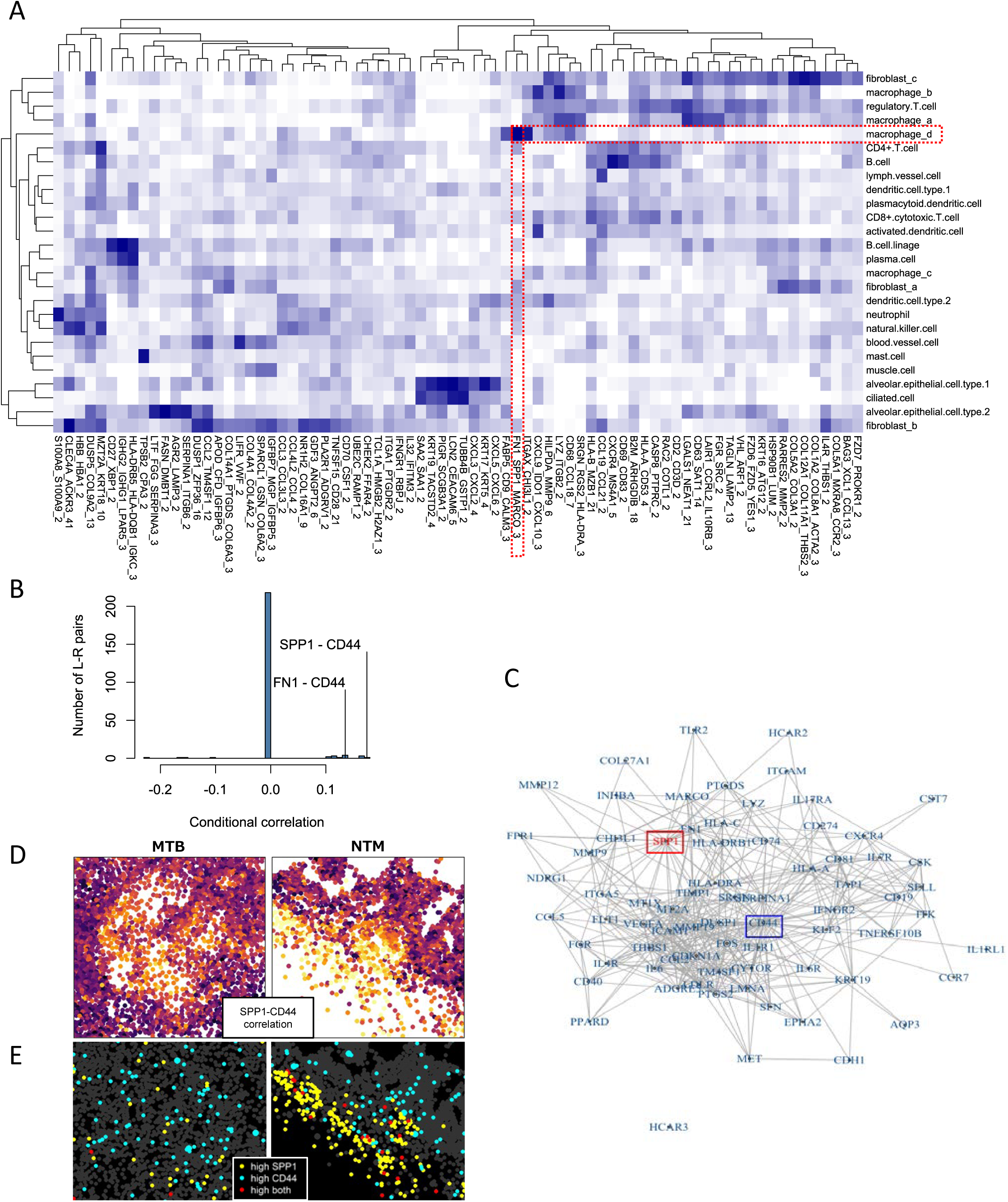
Coexpression and ligand-receptor assay in InSituCor. A. Heatmap of co-expressed transcripts across different cell types. Co-expression of FN1, SPP1, and MARCO was specifically observed in macrophage_d (highlighted by the broken red line). B. Conditional correlation of ligand-receptor pairs in the ligand-receptor assay. The SPP1-CD44 and FN1-CD44 pairs show strong conditional correlations. C. Correlation network of SPP1 and CD44- associated transcripts. D. Spatial images showing SPP1-CD44 correlations in MTB and NTM tissues. Yellow indicates high correlation in the images (left: MTB1, right: NTM2). E. Spatial images from CosMx™ analysis of MTB and NTM samples. High expression of SPP1, CD44, and both are indicated by yellow, blue, and red dots, respectively.

Dot plot analysis showed that CD44 was expressed by a variety of cell types, including macrophages, mast cells, CD8+ T cells, and dendritic cells around the granulomatous lesions, while SPP1 expression was predominantly confined to the macrophage_d subtype (Fig. 6A). Immunohistochemistry for SPP1 and CD44 showed SPP1 and CD44 expression in Langhans giant cells and various cells around SPP1-positive macrophages, respectively (Fig. 6B). Spatial imaging using the CosMx™ platform confirmed that CD44 was expressed not only in macrophages but also in various cell types, including B cells, T cells, and epithelial cells surrounding SPP1-positive macrophages (Fig. 6C). Immunofluorescence assay further confirmed that CD44 expression in SPP1-positive large foreign body macrophages and their surrounding cells (Fig. 6C). The SPP1-CD44 interaction was additionally identified using the CellChat package (Fig. 7). SPP1, expressed by macrophage_d, was shown to interact with various cell types. Circle plots and heatmaps of the SPP1 signaling pathway network indicated that the receiver cells for the SPP1-CD44 signal included macrophage_d, macrophage_c, and occasionally CD8+ cytotoxic T cells, while the sender cell was macrophage_d (SPP1-positive macrophage) (Figs. 7B, 7C, 7E, and 7G). The receiver cells of the SPP1-CD44 signaling pathway varied among samples (Fig. 7C). Although macrophages were the primary receiver cells, B cells, CD8+ T cells, and alveolar epithelial cells also acted as receivers in certain cases. Beyond the SPP1-CD44 signaling pathway, other significant cell communication pathways identified by CellChat included collagen, MHC-I, MIF, FN1, integrin β2, PECAM1, and ICAM pathways (Fig. 7D, fig. S11). The incoming and outgoing interactions of the SPP1-CD44 pathway by macrophage_d were relatively strong among the cell types and other pathways in mycobacterium-infected tissues (Figs. 7A, 7F, and 7G, fig. S11). In the SPP1-CD44 signaling pathway, SPP1 expressed by macrophage_d communicated with macrophage_d itself, mast cells, and CD8+ T cells (Figs. 7E-7G and fig. S11).

**Fig. 6.**
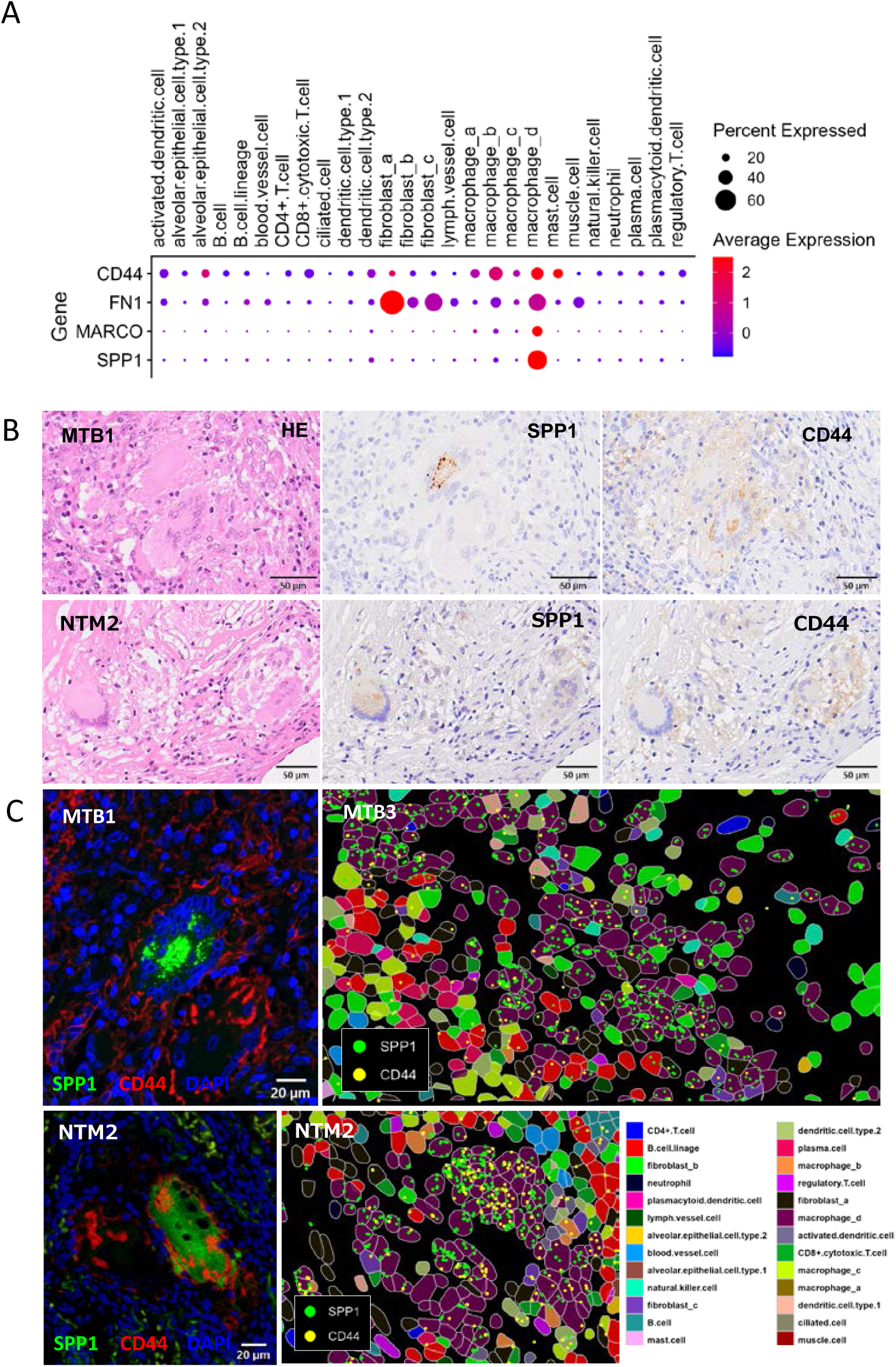
Expression of SPP1 and CD44 in macrophages around granuloma tissue with mycobacterial infection. A. Dot plot showing the expression of SPP1, MARCO, FN1, and CD44 across all cell types. B. Immunohistochemistry for SPP1 and CD44 in tissues from patients with MTB (upper panels) and NTM (lower panels) infections. C. Immunofluorescence and CosMx™ spatial images for SPP1 and CD44. SPP1 was expressed in the cytoplasm of multinuclear giant cells (immunofluorescence assay, left panels). CD44 expression was observed on the membrane of multinucleated giant cells and their neighboring cells. CosMx™ spatial images (right panels) showed that macrophage_d expresses both SPP1 and CD44. Colors representing different cell types are shown in the bottom right panel.

**Fig. 7.**
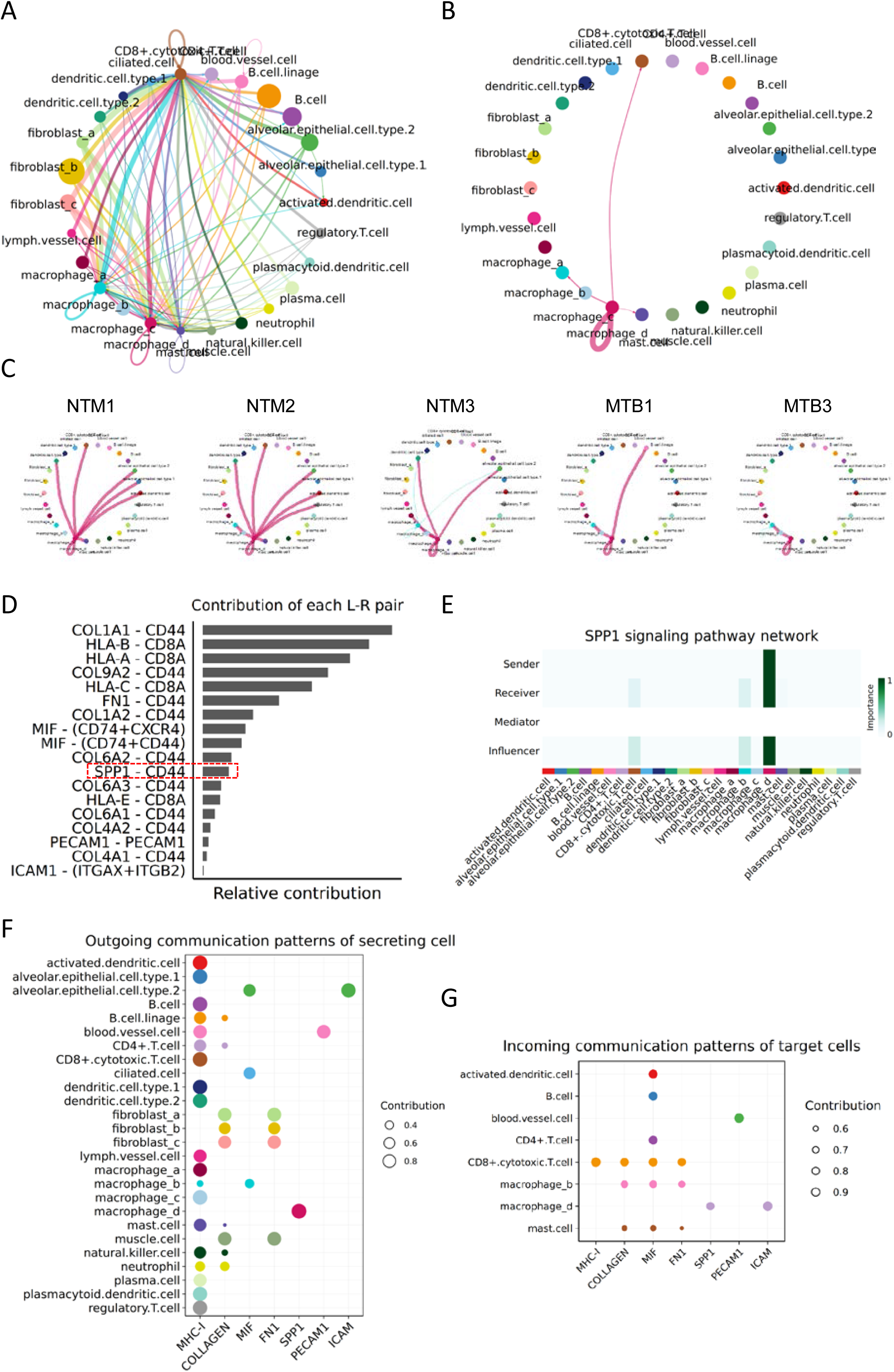
Analysis of cellular communication among cell types using CellChat. A and B. Circular plots showing interactions among cell types in all pathways (A) and specifically in the SPP1 signaling pathway (B) across all seven cases analyzed with CosMx™. C. Circular plots depicting interactions among cell types in the SPP1 signaling pathway in each individual case. D. Histogram showing the relative contribution of each ligand-receptor (L-R) pair across all seven cases analyzed with CosMx™. Only significant L-R pairs are shown, with SPP1 interacting exclusively with CD44. E. Heatmap illustrating the signal intensity of the sender and receiver in the SPP1 signaling pathway across all samples. F. Dot plots showing the outgoing communication patterns of secreting cells. Significant outgoing signals in the identified signaling pathways are represented by dots. G. Dot plots showing the incoming communication patterns of secreting cells. Significant incoming signals are represented by dots.

### SPP1-positive macrophages in granuloma associated with non-mycobacterial infection

To determine whether SPP1-positive macrophages are associated with granulomatous diseases beyond mycobacterial infections, pathological samples from granulomatous lesions in patients with rheumatoid nodule (RN), granulomatosis with polyangiitis (GPA), sarcoidosis (SA), Aspergillus pneumonia (AP), and aspiration pneumonia (Aspi-P) were examined. Immunohistochemistry revealed that SPP1 was present in some foreign body giant macrophages surrounding the granulomatous lesions across all tested samples (Fig. 8). RNA-seq analysis of RNA extracted from these granulomatous samples demonstrated elevated expression of SPP1 in the granulomatous lesions, similar to the expression levels observed in mycobacterial infections, compared with control lung and tonsil samples (fig. S7C). Expression of CD40L (CD154) and CD40 pair, another ligand-receptor pair reported as activation in granulomatous diseases(*15*), did not show significant difference between granulomatous samples and controls (Fig. 5B). Furthermore, immunohistochemistry revealed CD44 expression in SPP1-positive macrophages and neighboring cells within the lesions of GPA, SA, RN, and AP (Figs. 8A and 8B). Immunofluorescence imaging also demonstrated that a subset of SPP1-positive macrophages expressed CD44 (Fig. 8C). These findings suggest that SPP1-positive macrophages are broadly associated with granulomatous diseases, and that the SPP1-CD44 interaction is observed commonly in granulomatous pathology.

**Fig. 8.**
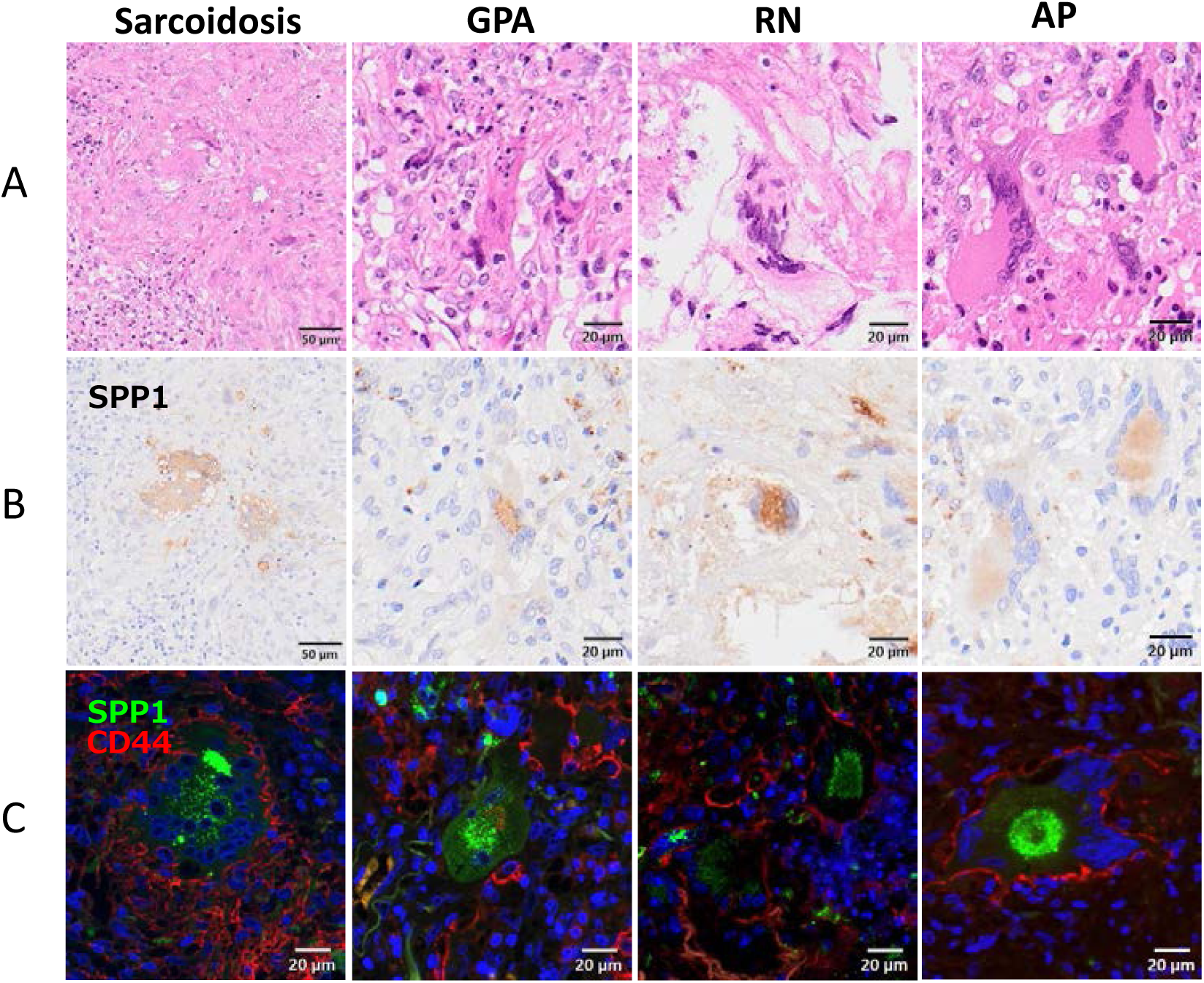
SPP1 and CD44 expression in tissues from patients with granulomatous diseases. A. HE staining. B. Immunohistochemistry for SPP1. C. Immunofluorescence of SPP1 and CD44. Tissues from patients with sarcoidosis (left), granulomatosis with polyangiitis (GPA, middle), and rheumatoid nodule (RN, right) were investigated.

## DISCUSSION

A pivotal discovery in our study was the identification of multiple macrophage subtypes within the tissues of granulomatous diseases. Based on gene expression patterns, we identified at least four distinct macrophage cell types. Among these, SPP1-positive macrophages predominantly infiltrated the regions surrounding NTM- and MTB-positive granulomatous tissue. Additionally, we found that Langhans giant cells were a specialized form of these SPP1-positive macrophages. Notably, besides these giant cells, we observed a significant number of SPP1-positive macrophages that did not exhibit giant cell morphology, indicating a diverse range of morphologies for these cells.

Our spatial transcriptomics data demonstrated that at least 26 types of cells infiltrated the lungs with mycobacterial infection (Fig.1). Although the percentages of infiltrating cell types varied among samples, macrophages were predominant cell types adjacent to granuloma regions in all seven samples. Along with macrophages, T cells, B cells, plasma cells, and dendritic cells were also observed around granuloma. These cell populations are consistent with those reported in MTB and NTM tissues in previous studies (*2–9*). Macrophages have been shown to be key player in granuloma formation (*5, 7, 10*). Various macrophage subtypes infiltrate granulomas alongside T and B cells in mycobacterial infections, with M1/M2 macrophage polarization inducing NF-kappa B activation, which is associated with granuloma formation and outcome(*10*).

SPP1, also known as osteopontin, is a multifunctional secreted phosphorylated glycoprotein produced by macrophages and T cells. It is associated with bone remodeling, T cell and B cell activation, and chemotaxis(*16–20*). SPP1 expression in foreign body macrophages, such as Langhans giant cells, has been previously reported in pathological samples from tuberculosis and sarcoidosis (*16, 21–24*). However, there are no reports describing macrophages without large cell morphology expressing SPP1 around granulomas. Given that SPP1-positive macrophages share a similar transcript profile with Langhans giant cells and are categorized within the same cluster (Fig. 3), some of these cells may subsequently fuse to form Langhans giant cells. Recent studies using single cell transcriptomics and/or spatial omics assay have highlighted the potential role of SPP1-positive macrophages in infections, inflammatory diseases and cancer (*25, 26*). For instance, in tissue samples from a patient with granulomatous slack skin, CD11c+/LYZ+ macrophages exhibited an M1-like phenotype and strongly expressed MMP9, MMP12, CHI3L1, CHIT1, COL1A1, TIMP1, and SPP1, all of which are involved in extracellular matrix degradation and tissue remodeling (*27*). SPP1-positive macrophages have also been detected in bronchoalveolar lavage fluid from tuberculosis patients (*28*). Interestingly, plasma osteopontin levels are higher in patients with tuberculosis and sarcoidosis compared to healthy donors (*29, 30*). A study on perivascular adipose tissue in heart transplant recipients demonstrated the presence of SPP1-positive macrophages in the perivascular tissues, with an identified interaction between secreted SPP1 and CD44 (*31*). These data suggest that SPP1-positive macrophages secrete SPP1 and induce SPP1-CD44 signaling in inflammation. In a *Schistosoma mansoni* egg-induced granuloma animal model, granulomas from osteopontin-knockout animals were smaller and contained significantly fewer macrophages, as well as fewer macrophage-derived epithelioid cells and giant cells (*32*). Thus, the presence of SPP1-positive macrophages in granulomatous tissues suggests an important role for SPP1 in granuloma formation through its interactions.

In our study, we demonstrated that the SPP1-CD44 pathway was one of the main ligand-receptor pairs involved in cell-cell communication in granulomatous lesions (Fig. 7). Recent studies have shown that SPP1 is associated with cancer cell growth and resistance to chemoradiotherapy, inducing epithelial-mesenchymal transition, autophagy, aberrant glucose metabolism, epigenetic alterations, and reduced drug uptake through the SPP1-CD44 signaling (*33*). SPP1-positive macrophages have been identified in colorectal cancer and squamous cell carcinoma(*34, 35*). In hepatocellular cancer, SPP1-positive macrophages infiltrated hypoxic tumor regions with cancer stem cells, where the SPP1 signaling pathway was highly active in cell-cell communication with SPP1-CD44 and SPP1-ITGA/ITGB identified as the main ligand-receptor pairs (*36*). SPP1 also binds to integrins α4β1, α9β1, and α9β4, inducing cell adhesion and migration. CD44 has also been identified as a receptor for SPP1, and the SPP1-CD44 signaling pathway has been reported in various cancers(*37–39*). SPP1-positive macrophages have been detected in cancers as tumor-associated macrophages, and the SPP1-CD44 pathway has been identified as a critical mediator of communication between tumor-associated macrophage subpopulations (*40–42*). The SPP1-CD44 pathway has been linked to cancer chemoresistance in solid tumors and to cell-to-cell communication between cancer cells and tumor-associated macrophages (*36, 43, 44*). The presence of SPP1-positive macrophages in cancers correlates with worse prognosis and poor immunotherapy response, leading to reduced survival (*36, 40*). Some therapeutic approaches targeting SPP1-CD44 signaling have been reported in vitro and in vivo. In a mouse model, SPP1 was shown to induce cancer cell chemoresistance by activating the CD44 receptor, and that treatment with anti-SPP1 and anti-CD44 antibodies improved cancer cell sensitivity to cisplatin (*45*). Importantly, the SPP1-CD44 signaling pathway has been identified as a potential immune checkpoint target(*46, 47*). CD44 expression in tumor cells was positively correlated with PD-L1 expression and the numbers of infiltrated CD68- and CD168-positive macrophages(*48*). It was also demonstrated that SPP1-CD44 immune checkpoint regulates CD8+ T cell activation and tumor immune evasion in interferon regulatory factor 8 deficient mice (*46*). Inhibition of the SPP1-CD44 pathway, therefore, may be a potential therapeutic strategy to reduce granuloma formation in granulomatous diseases, such as mycobacterial infections. In tuberculosis and MAC infections, granulomas encapsulate mycobacteria in necrotic tissue to prevent their spread. Since granuloma formation is a major characteristic of mycobacterial infections, further studies will be required to explore the inhibition of the SPP1-CD44 pathway in macrophages.

In conclusion, our study sheds light on the intricate cellular dynamics involved in granuloma formation in mycobacterial diseases. The prominence of SPP1-positive macrophages, including Langhans giant cells, suggests that SPP1-CD44 pathway in these cells may play a role in this process, opening avenues for further research and potential therapeutic interventions.

## MATERIALS AND METHODS

### Tissue samples

FFPE samples of resected lung tissues from patients with active mycobacterial diseases, granulomatous diseases, or other controls were used (table S1). For CosMx™ spatial molecular imaging (SMI) analysis(*14*), 7 lung resections were obtained from 3 MTB and 4 NTM patients undergoing lung resection surgery. Total 28 lung tissues from 25 patients were applied to histological analysis, immunohistochemistry and RNAseq analysis by next generation sequencing. In addition, five intact lung tissues from patients with cancer and four tonsillar tissues were used as controls. All MTB and NTM were diagnosed as active disease by sputum culture and/or PCR method. The study was approved by the Ethics Committee of National Institute of Infectious Diseases (approval no. 1781) and National Hospital Organization Tokyo National Hospital (approval no. 408).

### Single cell spatial transcriptomics

Spatial transcriptomic profiling of FFPE lung sections was performed using the CosMx^TM^ Spatial Molecular Imager, as previously described (*14*). The CosMx™ Human Universal Cell Characterization RNA Panel (1000-plex; Nanostring, Seattle, WA) was applied to seven FFPE samples, including three tuberculosis samples, three MAC-positive pneumonia samples, and one MAC-negative bronchiectasis sample with 15-45 fields of view (fov) per sample, totaling 216 fovs (Table S3). Fovs measuring 0.512 mm × 0.512 mm were used for the commercial instrument in the final runs (MTB2, MTB3, NTM3), whereas fovs measuring 0.985 mm × 0.657 mm were used for the development instrument in the remaining samples (MTB1, NTM1, NTM2, NTM4). Fovs were placed on the tissue to align with regions of interest identified by H&E staining of an adjacent serial section. Fovs were selected from areas around granuloma or inflammatory cell infiltration to avoid granulomatous necrotic tissues. The mean area explored was 4.60 mm^2^ per sample. Sample preparation and CosMx™ SMI instrument run was performed as described previously (*14, 25*). In addition to RNA readout, tissue samples were incubated with a fluorophore-conjugated antibody cocktail against CD298/B2M (488 nm), PanCK (532 nm), CD45 (594 nm), and either CD3 or CD68 (647 nm) proteins and DAPI stain. Cell segmentation, assigning transcripts at the cell-level, and obtaining a transcript by cell count matrix were performed as previously described (*14, 25*). Following total count normalization across all detected cells, highly variable genes (HVGs) were identified. Then, principal component analysis (PCA) was computed on the HVGs. Finally, uniform manifold approximation and projection (UMAP) was performed on these data(*14, 25*).

### Cell type annotation

To identify subpopulation markers, we ran findMarkers (scran package, version 1.22.1)(*49*), blocking by sample, considering HVGs only, and testing for positive log-fold changes. The 100 top ranked genes were selected and passed to SingleR (version 1.8.1)(*50*).

### Co-expression and cellular communication assay

Spatial correlation analysis such as co-expression and receptor-ligand assay was performed in the CosMx™ dataset with a R package InSituCor (*51*). InSituCor was run on normalized count data, adjusted for the effects of cell type, tissue ID and single cell counts, to uncover spatial correlations beyond those accounted for by these variables. Cellular communication for cell types were also analyzed with CellChat (http://www.cellchat.org/). In CellChat, to convert spatial transcriptomics data of CosMx™, conversion.factor = 0.18 was used for setting spatial factors. In function “computeCommunProb”, contact.range = 50 and scale.distance = 20 were used.

### Bulk RNA-seq analysis of lung tissues

Total RNA was extracted from FFPE samples (Supplementary Table 1, 5 MTB, 7 NTM, 12 granulomatous diseases including 2 RN, 3 AP, 3 GPA, 3 SA, and 1 Aspi-P, 4 control lung, and 4 tonsils) with Quick-DNA/RNA FFPE Miniprep Kit (Zymo Research). Barcoded RNA-seq library was established with NEBNext Ultra II RNA Library Prep Kit for Illumina (New England BioLabs, Ipswich, MA) according to the manufacturer’s instructions. Libraries were subjected to pair-end sequencing (300 cycles) on a Nextseq 1000 platform (Illumina, San Diego, CA). Quality filtering, adapter trimming, and RNAseq analyzing was performed using CLC genomics workbench (version 23, Qiagen, Hilden, Germany). Homo sapience refseq GRCh38.p13 was used as a reference genome of RNA sequences. PCA, volcano blot, and box plots were performed using the R (version 4.3.1) statistics package with RStudio (version 2023.06.1+524). Gene ontology assay was performed in biojupies (https://maayanlab.cloud/biojupies/analyze/).

### Immunohistochemistry (IHC), immunofluorescence (IF), and special staining

Immunostaining was performed in the tissue sections of mycobacterial infection and other granulomatous diseases. Anti-SPP1 rabbit polyclonal antibody (HPA027541, Sigma-Aldrich, St. Louis, MO) and anti-CD44 mouse monoclonal antibody (UMAB134, OriGene Technologies, Rockville, MD) were used as primary antibodies. Anti-Bacillus Calmette– Guérin rabbit polyclonal antibody (B124, Dako, Copenhagen, Denmark) was used for detecting mycobacterial infection. For the second and third phase immunostaining reagents, a biotinylated F(ab’)2 fragment of rabbit anti-mouse immunoglobulin (Dako) and peroxidase-conjugated streptavidin (Dako) were used. 3-3’diaminobenzidine was used as a chromogen and the slides were counterstained with hematoxylin. Nuclear counterstaining was performed with DAPI on IF samples. Image acquisition was performed on virtual slide system (VS200, Evident, Tokyo, Japan). The Ziehl-Neelsen stain was also performed to confirm mycobacterial infection.

## Acknowledgments

The authors thank Mike McKenna, Claire Williams, Patrick Danaher, Joseph M. Beechem, Nanostring USA, Fumiyo Kurokawa, Takayuki Taniya, and Takashi Uematsu, Nanostring Japan, and Ikuyo Yamamoto, Department of Pathology, National Institute of Infectious Diseases, for technical supports. The authors also thank Akiko Yamashita, Yukari Nogi, and Ginko Kaneda, Department of Mycobacteriology, National Institute of Infectious Diseases, for their assistance.

## Author contributions

Conceptualization: HK, YH

Methodology: HK, YS

Investigation: HK, AH, YS, YH

Visualization: HK, YH

Funding acquisition: HK, YH

Project administration: HK, YH

Supervision: HK, YH

Writing – original draft: HK, YH

Writing – review & editing: HK, AH, YS, YH

## Data and materials availability

Bulk RNA-seq and CosMx™ SMI raw data generated in this study have been deposited in the GEO Omnibus database under accession code GSE276060 and GSE276083, respectively. The Seurat and Giotto objects of spatial analysis by CosMx™ SMI are registered in Figshare (https://figshare.com/). Full code for analysis of CosMx™ SMI data can be found at https://github.com/katanoh/Lung_CosMx_analysis.

## Supplementary Infomation

**Supplementary Fig.S1a:**
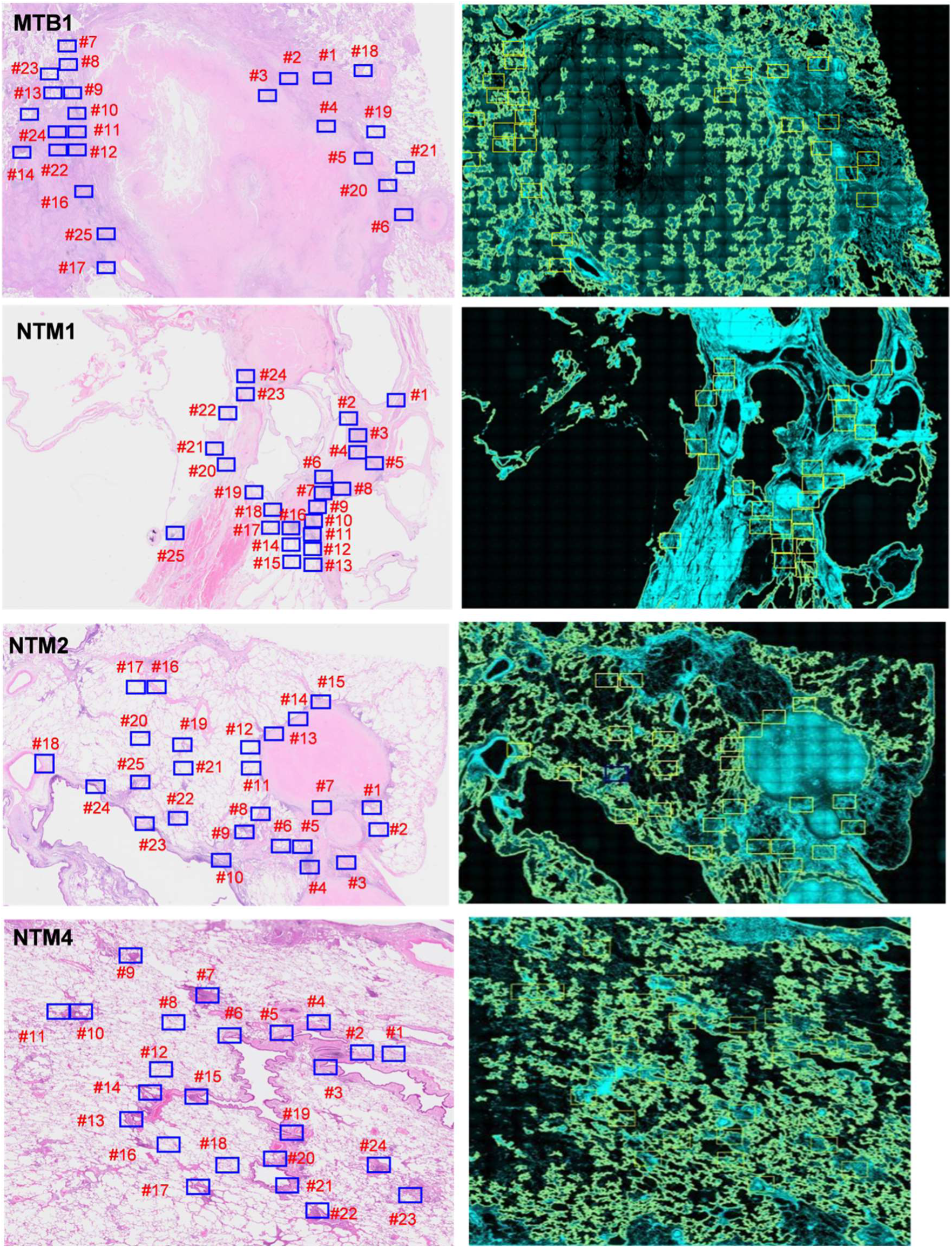
CosMx FOV placements.

**Supplementary Fig.S1b:**
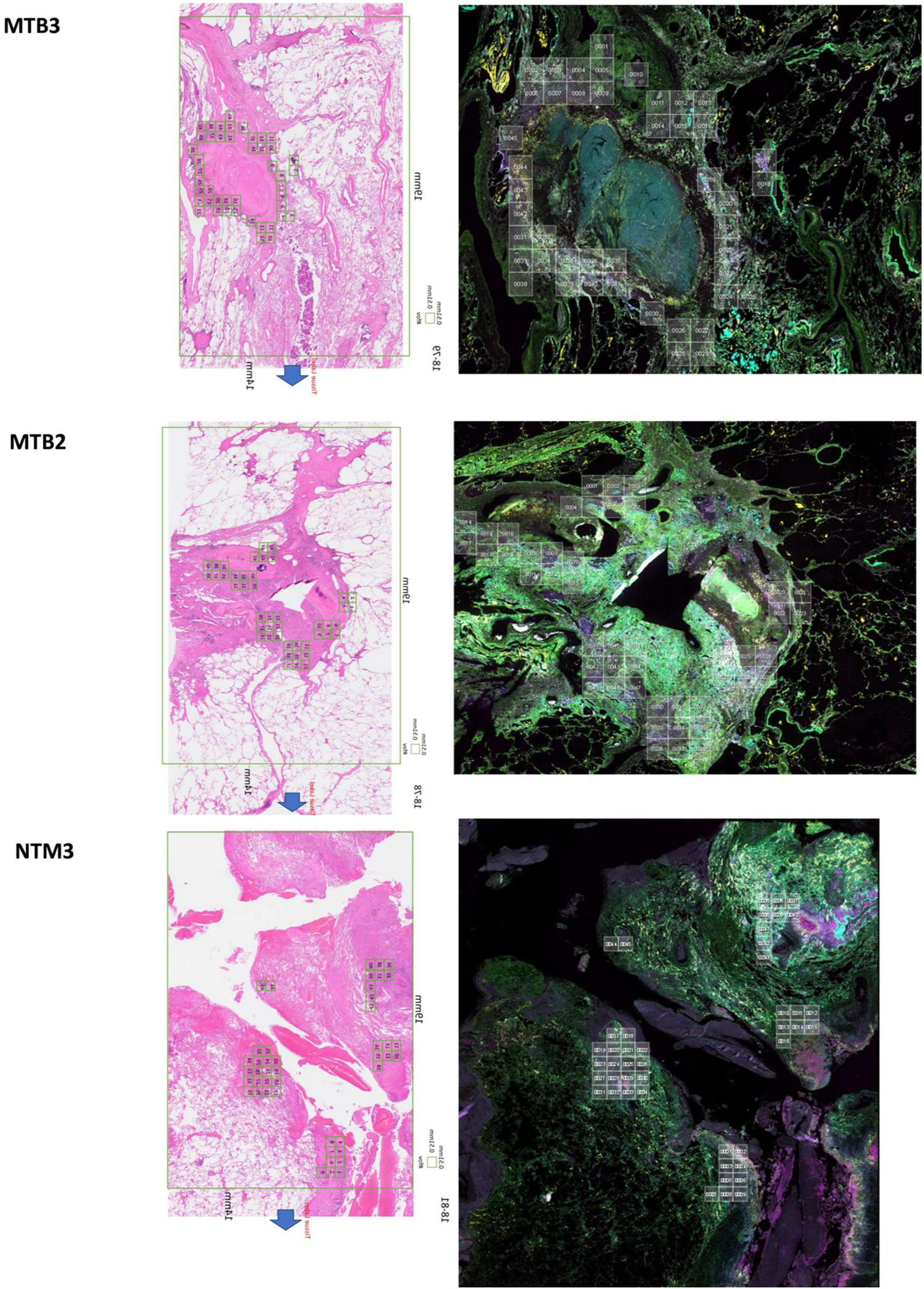
CosMx FOV placements.

**Supplementary Fig. S2:**
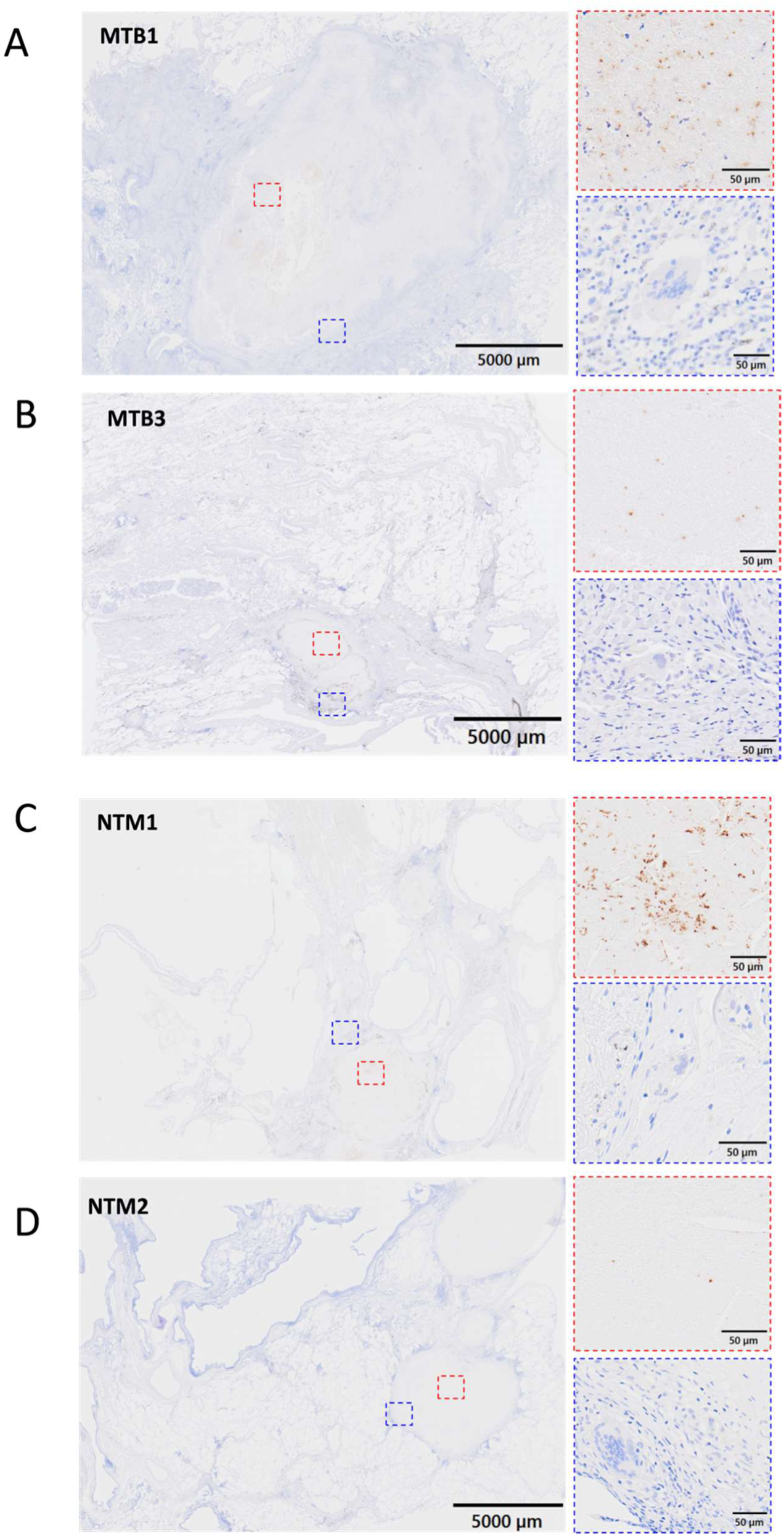
Immunohistochemistry for BCG. Blue and red boxes (right panels) show the enlarged images of left panels.

**Supplementary Fig. S3:**
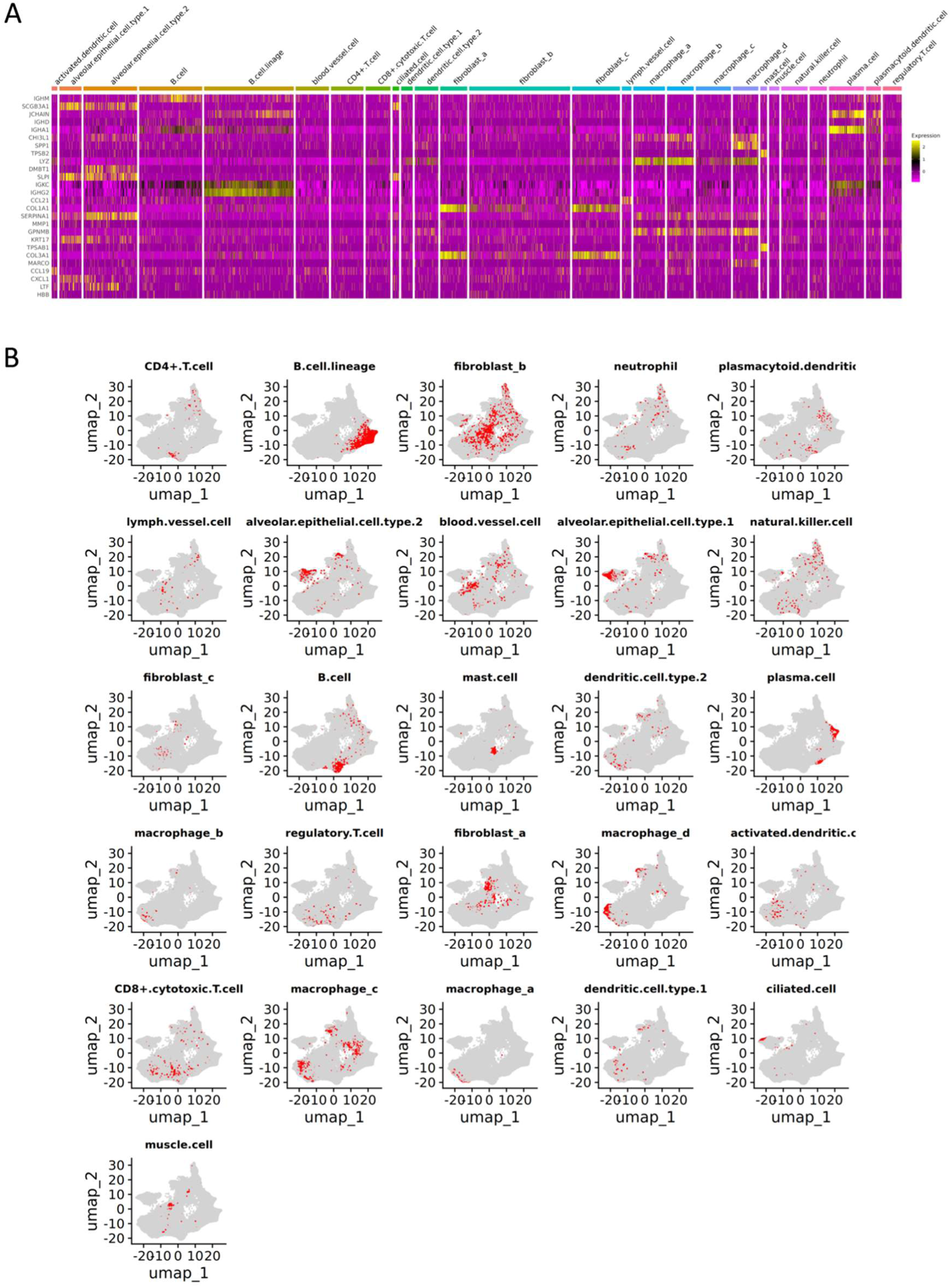
Single cell analysis of CosMx™ in mycobacterium-infected tissues. A. Heatmap showing gene expression and cell types based on single-cell data from CosMx™. B. UMAP visualization of cell types, with each cell type displayed in a red color.

**Supplementary Fig. S4:**
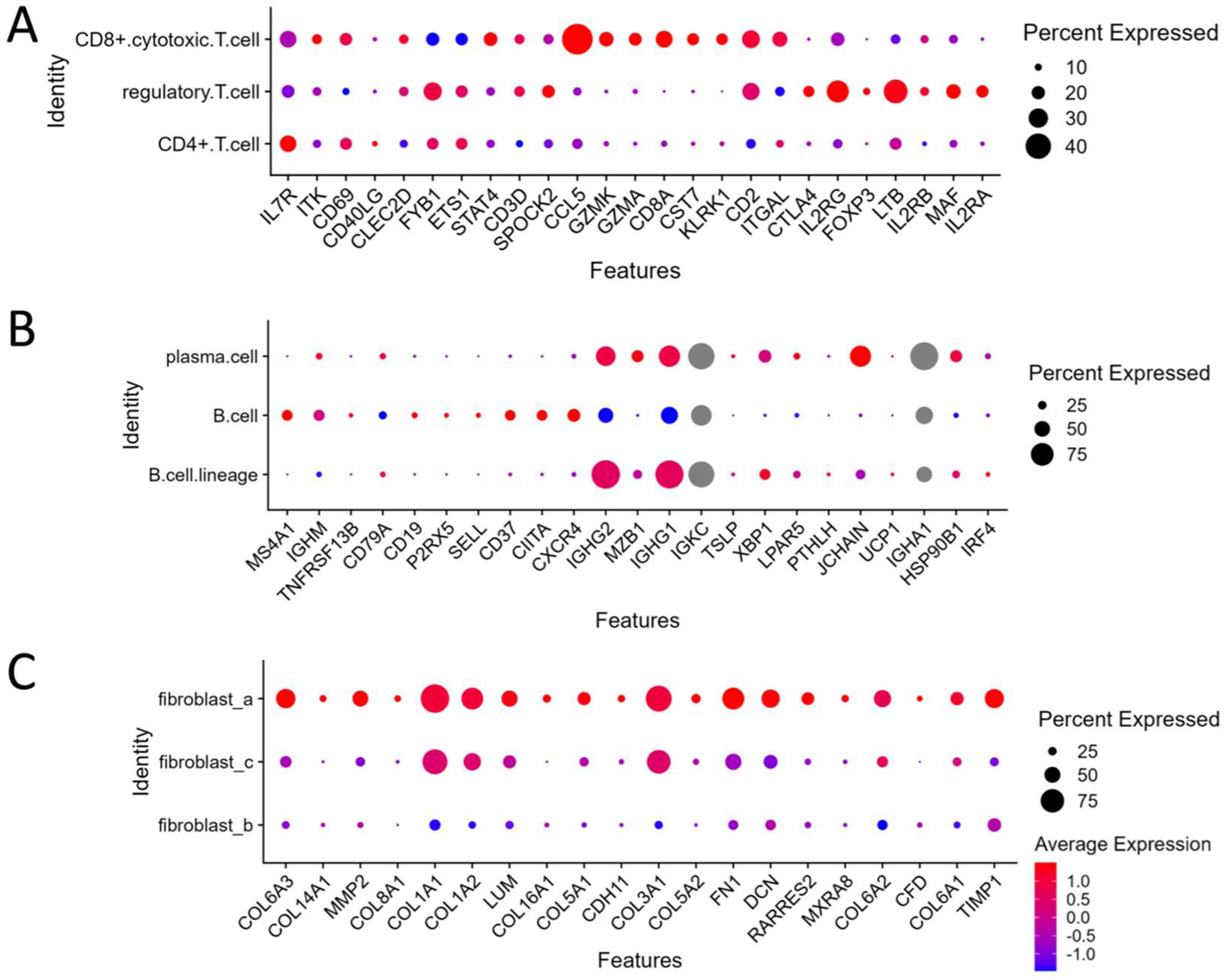
Dot plot showing gene expression in subtypes of T cells, B cells, and fibroblasts in single-cell analysis of CosMx™ in mycobacterium-infected tissues. A. Dot plot displaying gene expression in three T cell subtypes. B. Dot plot displaying gene expression in three B cell subtypes. C. Dot plot displaying gene expression in three fibroblast subtypes.

**Supplementary Fig. S5:**
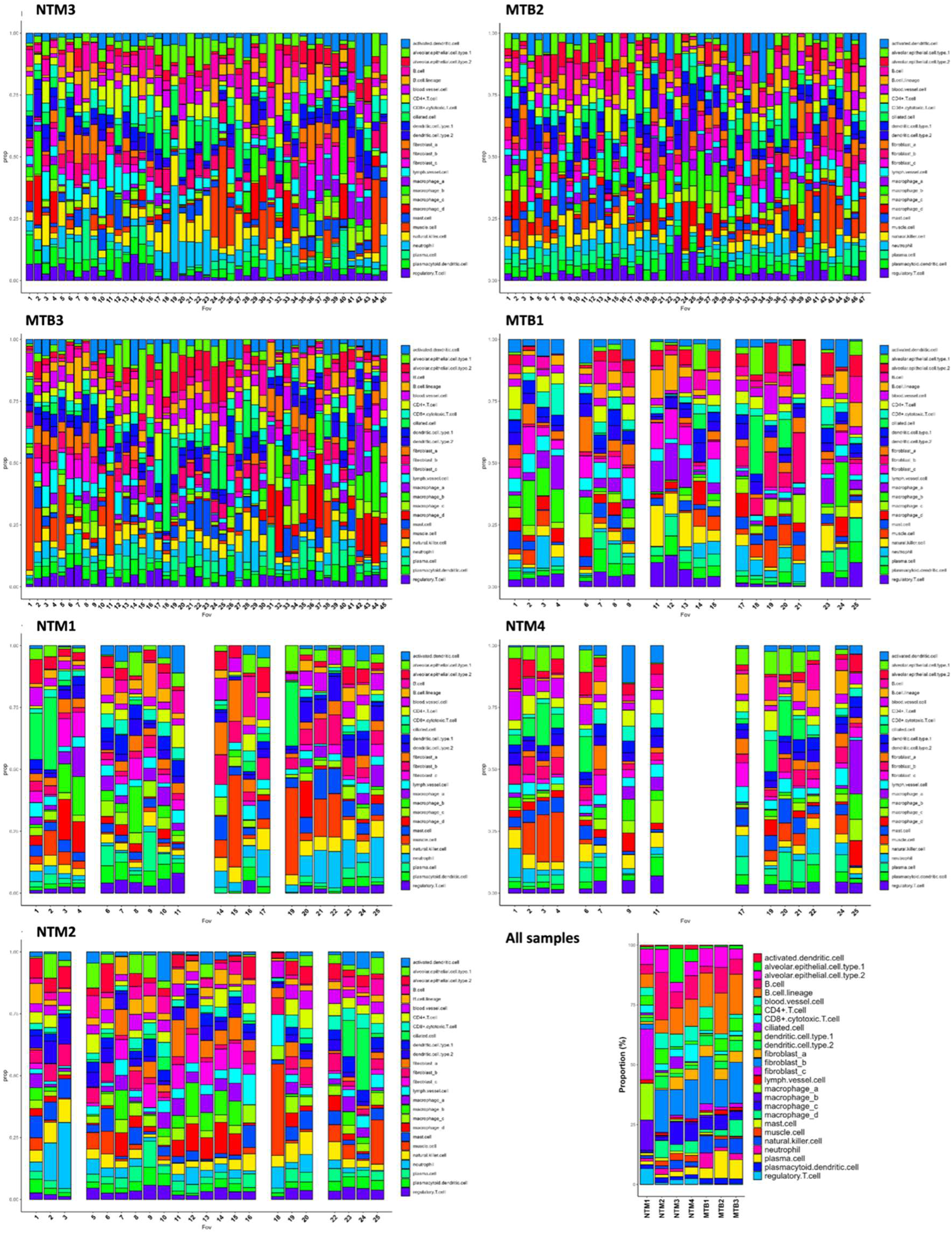
Populations of cell types in each tissue. Distribution of cell type across fovs in each slide and all samples (the right bottom panel).

**Supplementary Fig. S6a:**
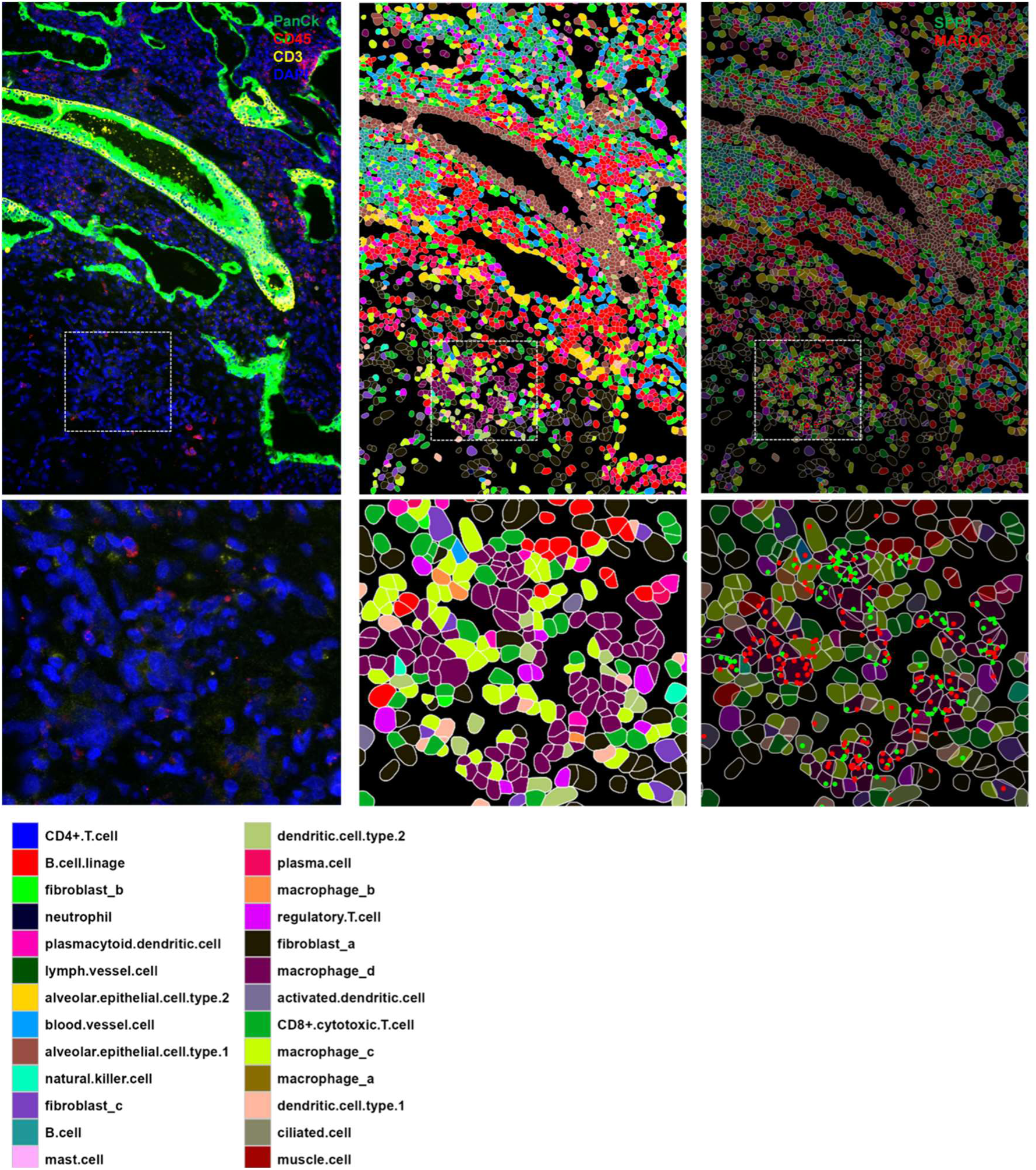
CosMx™ spatial images of SPP1 and MARCO expression in macrophage_d within MAC+ tissue (NTM2 fov9). Upper panels: Macrophage_d in the interstitial region expressing SPP1 and MARCO. Lower panels: Enlarged view of upper panel. Colors indicating cell types are shown in the bottom panel.

**Supplementary Fig. S6b:**
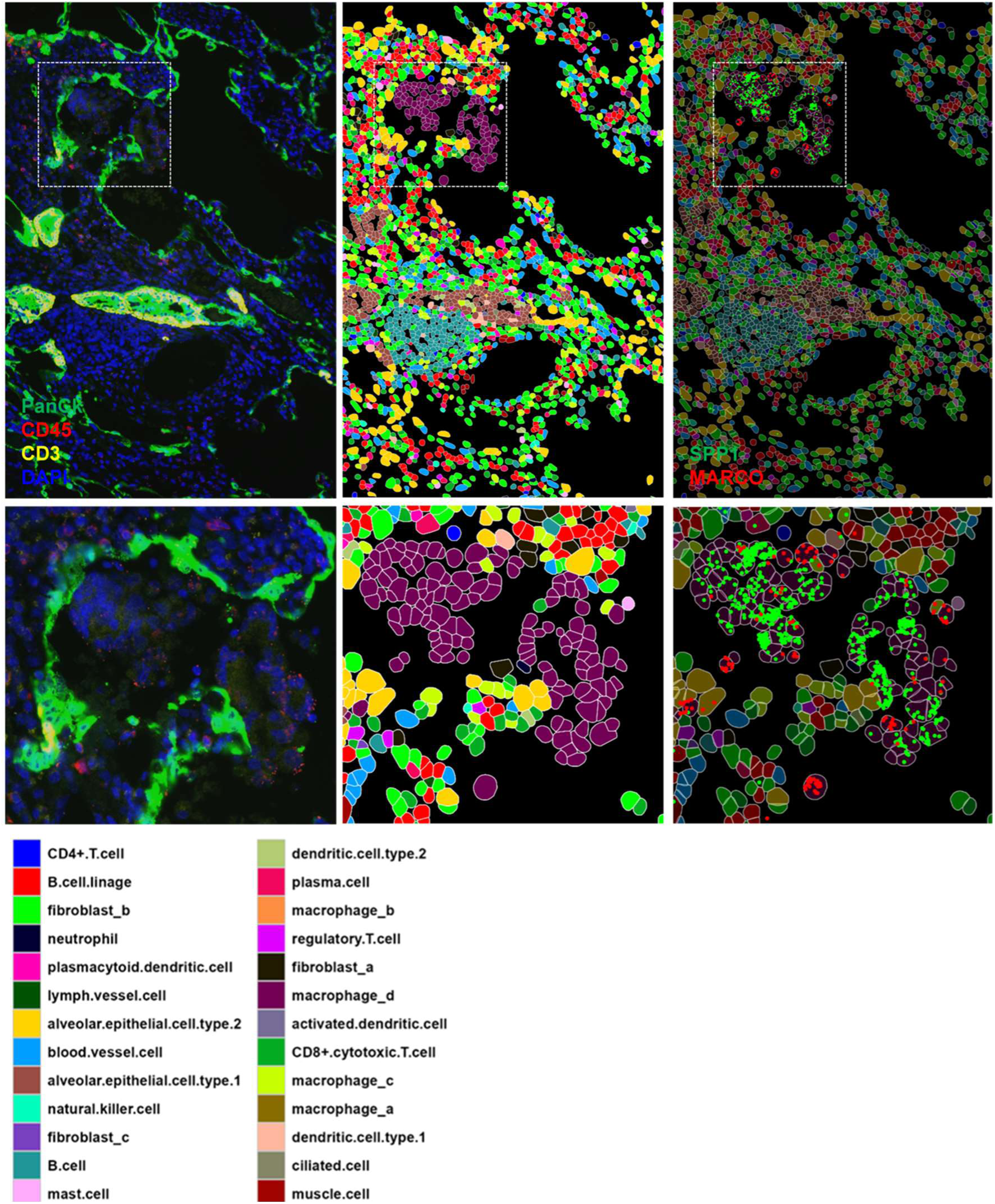
CosMx™ spatial images of SPP1 and MARCO expression in macrophage_d within MAC+ tissue (NTM2 fov8). Upper panels: Macrophage_d in the interstitial region expressing SPP1 and MARCO. Lower panels: Enlarged view of upper panel. Colors indicating cell types are shown in the bottom panel.

**Supplementary Fig. S6c:**
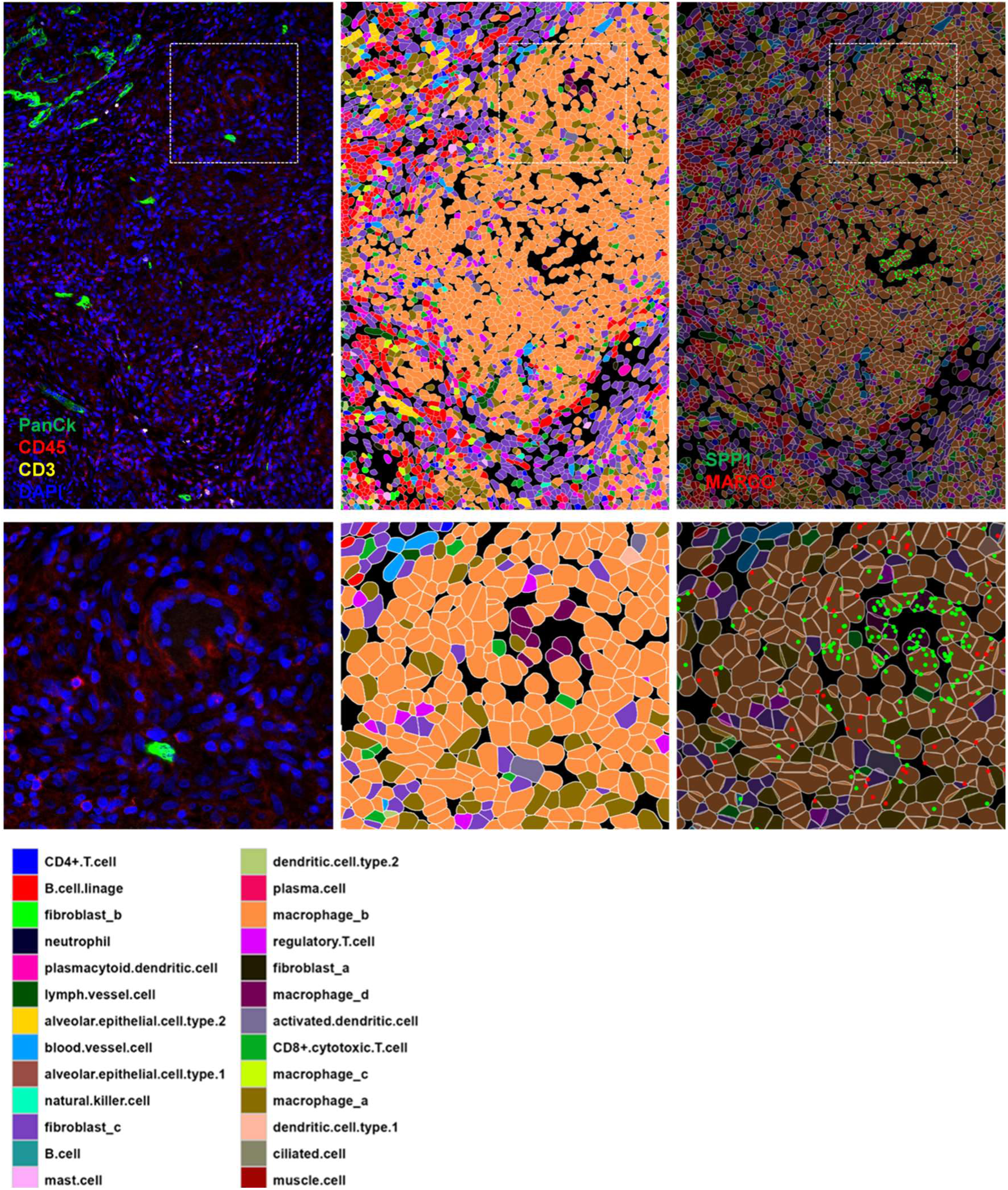
CosMx™ spatial images of SPP1 and MARCO expression in macrophage_d within MTB+ tissue (MTB1 fov24). Upper panels: Macrophage_d in the interstitial region expressing SPP1 and MARCO. Lower panels: Enlarged view of upper panel. Colors indicating cell types are shown in the bottom panel.

**Supplementary Fig. S7:**
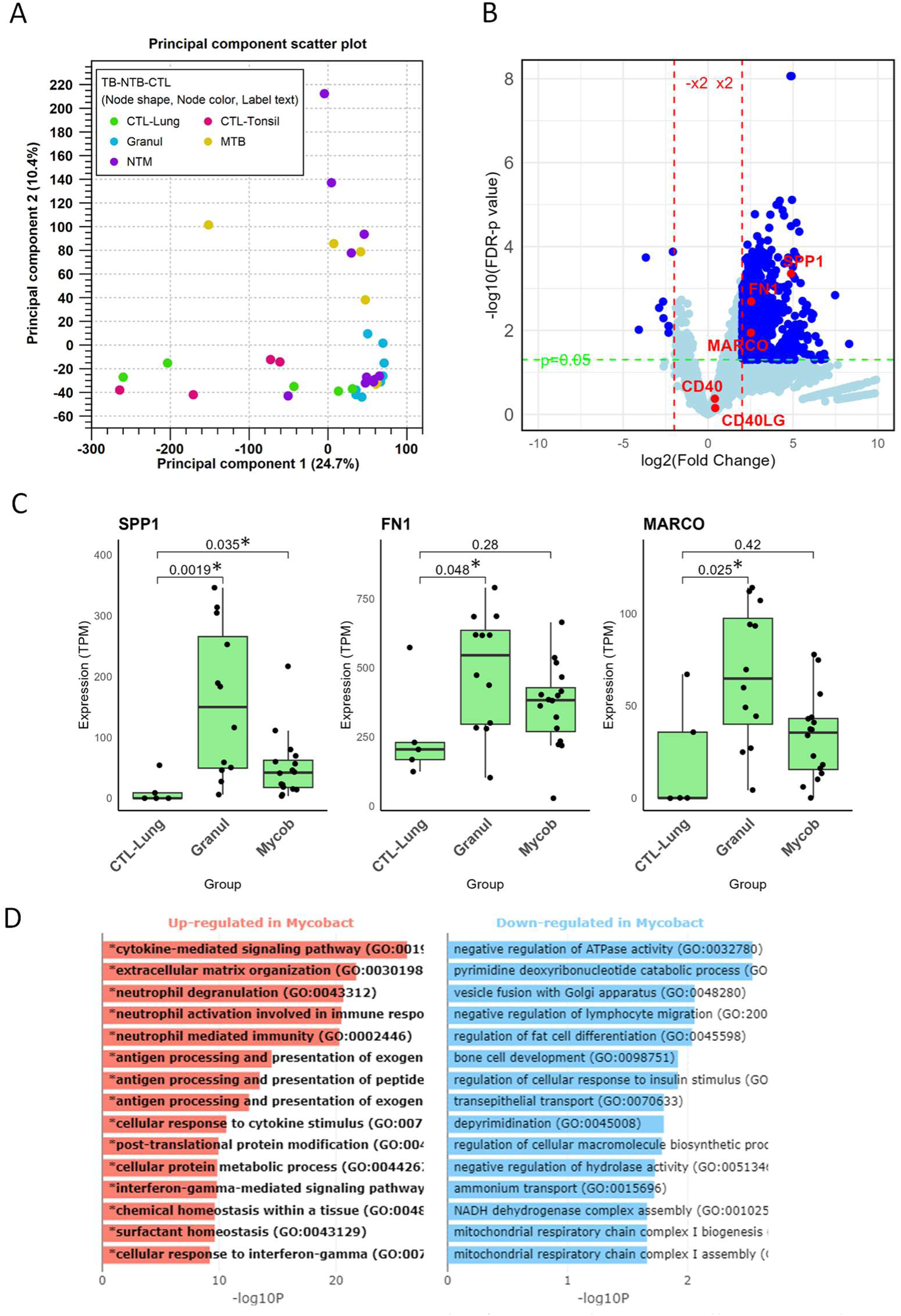
RNAseq Results for granulomatous diseases and mycobacterial infection. A. Principal component analysis (PCA) of RNA-seq data from 37 samples. B. Volcano plot comparing all granulomatous diseases (mycobacterial infections and granulomatous diseases other than mycobacterial infections) to control groups. The expression levels of SPP1, FN1, and MARCO are significantly higher in the overall granulomatous disease group. C. Box plot showing SPP1, FN1, and MARCO expression in control lung (CTL-Lung, n=5), granulomatous diseases other than mycobacterial infection (Granul, n=12), and the mycobacterial infection group (MTB+NTM, Mycob, n=16). Transcripts per million (TPM) values for each gene are plotted. P-values are displayed on comparison lines, with asterisks (*) indicating significant differences (Mann-Whitney test). D. Gene Ontology (GO) analysis comparing mycobacterial infection tissues to control lungs, with upregulated pathways listed in the left panel.

**Supplementary Fig. S8:**
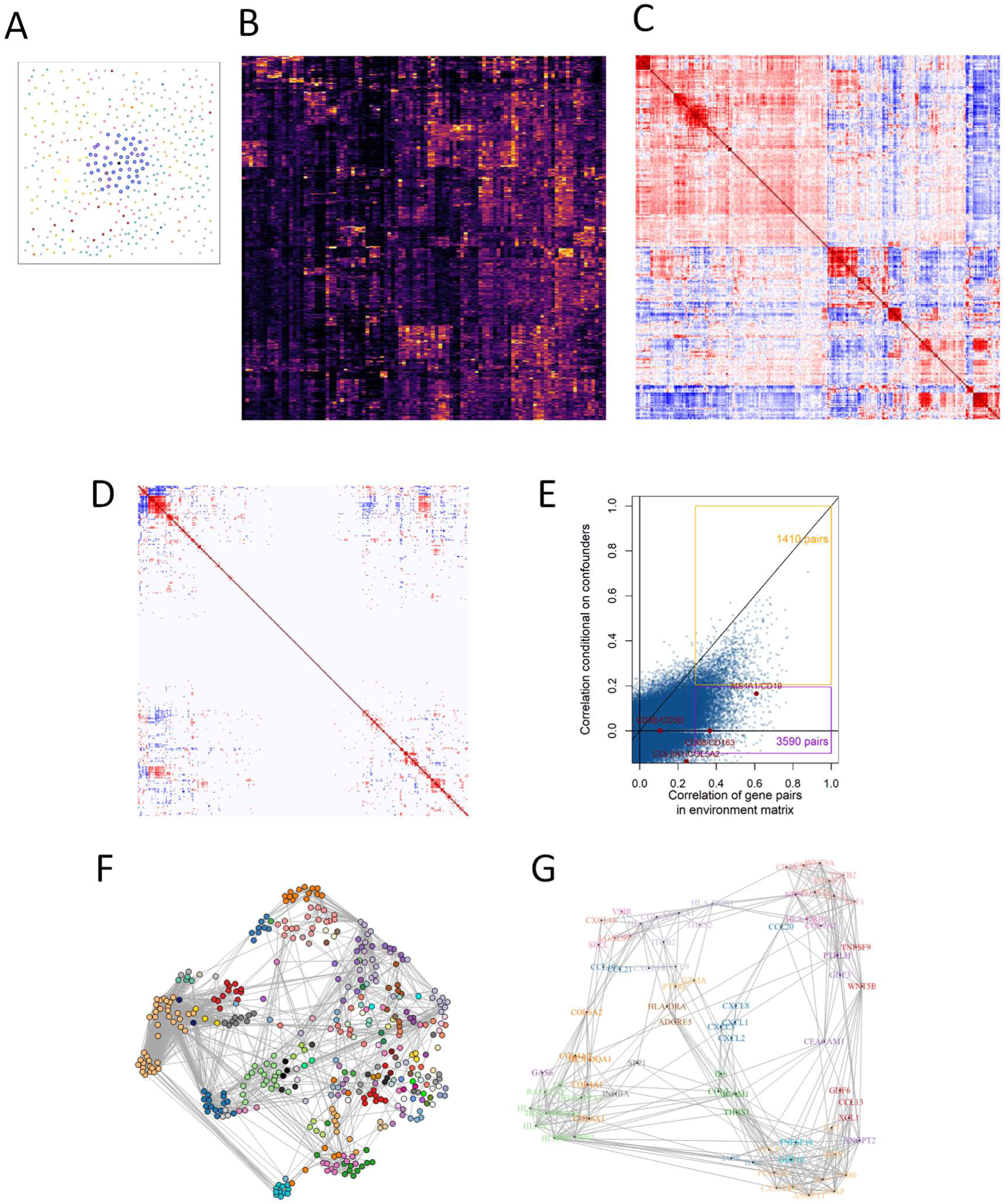
InSituCor analysis in MTB and NTM samples. A. Example of a cell’s nearest neighbors, used to define its “environment expression” profile. B. Subset of the environment expression matrix. C. Raw correlation matrix of the environment expression matrix showing near-ubiquitous correlations (subset of 300 random genes). D. Correlation matrix of the environment expression matrix (B), conditioned on the confounding matrix, over the same subset of genes as in (C). E. Raw vs. conditional correlation of environment gene expression. Selected pairs of marker genes are highlighted. F. Network representation of correlation between all genes in all modules. Genes with correlation > 0.2 are connected. G. Correlation structure of 51 ligands assigned to modules. Edges show conditional correlations > 0.1, and color shows module membership.

**Supplementary Fig. 9.**
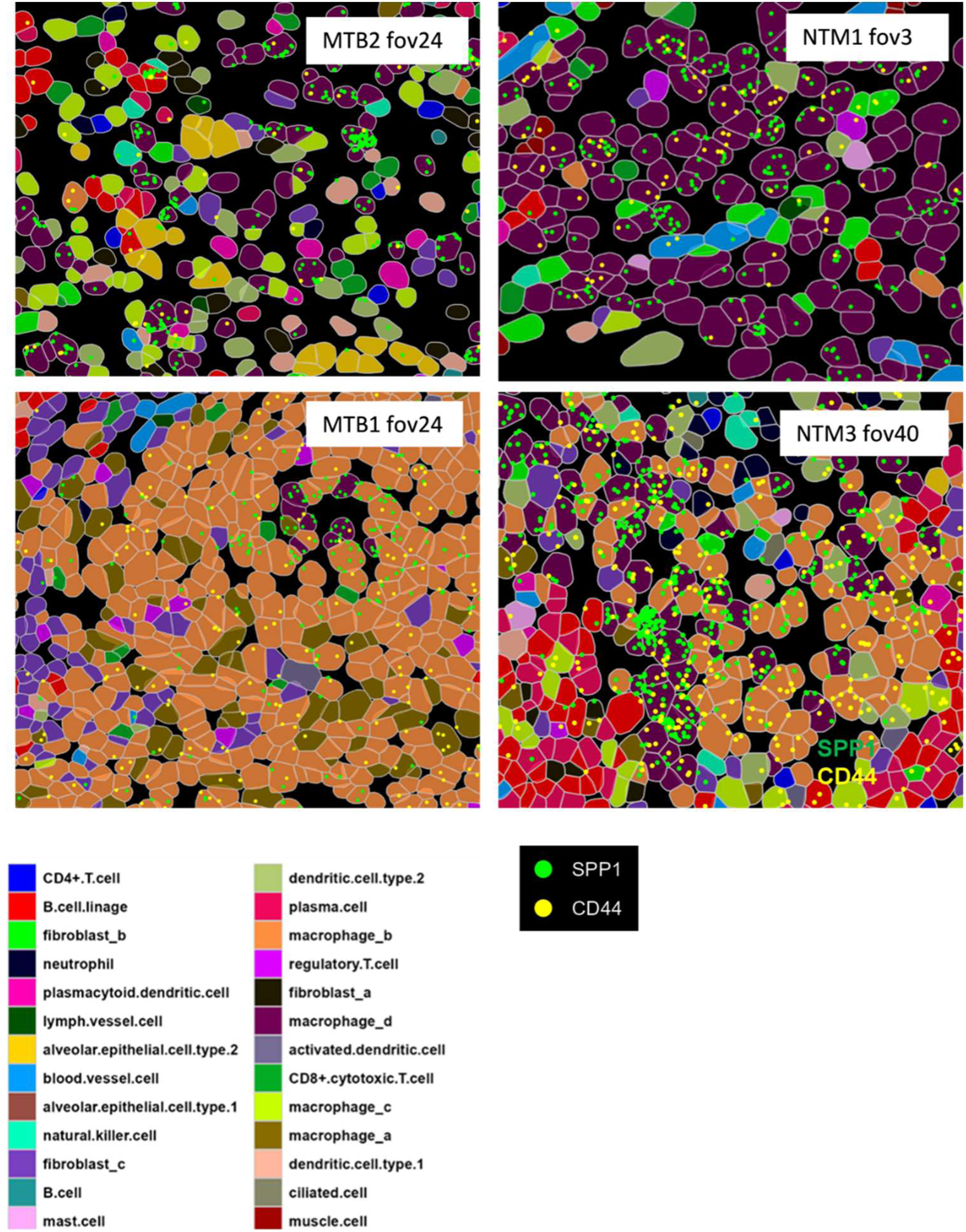
CosMx spatial images of SPP1 and CD44 expression in MTB and NTM tissues. Colors indicating cell types are shown in the bottom panel.

**Supplementary Fig. S10.**
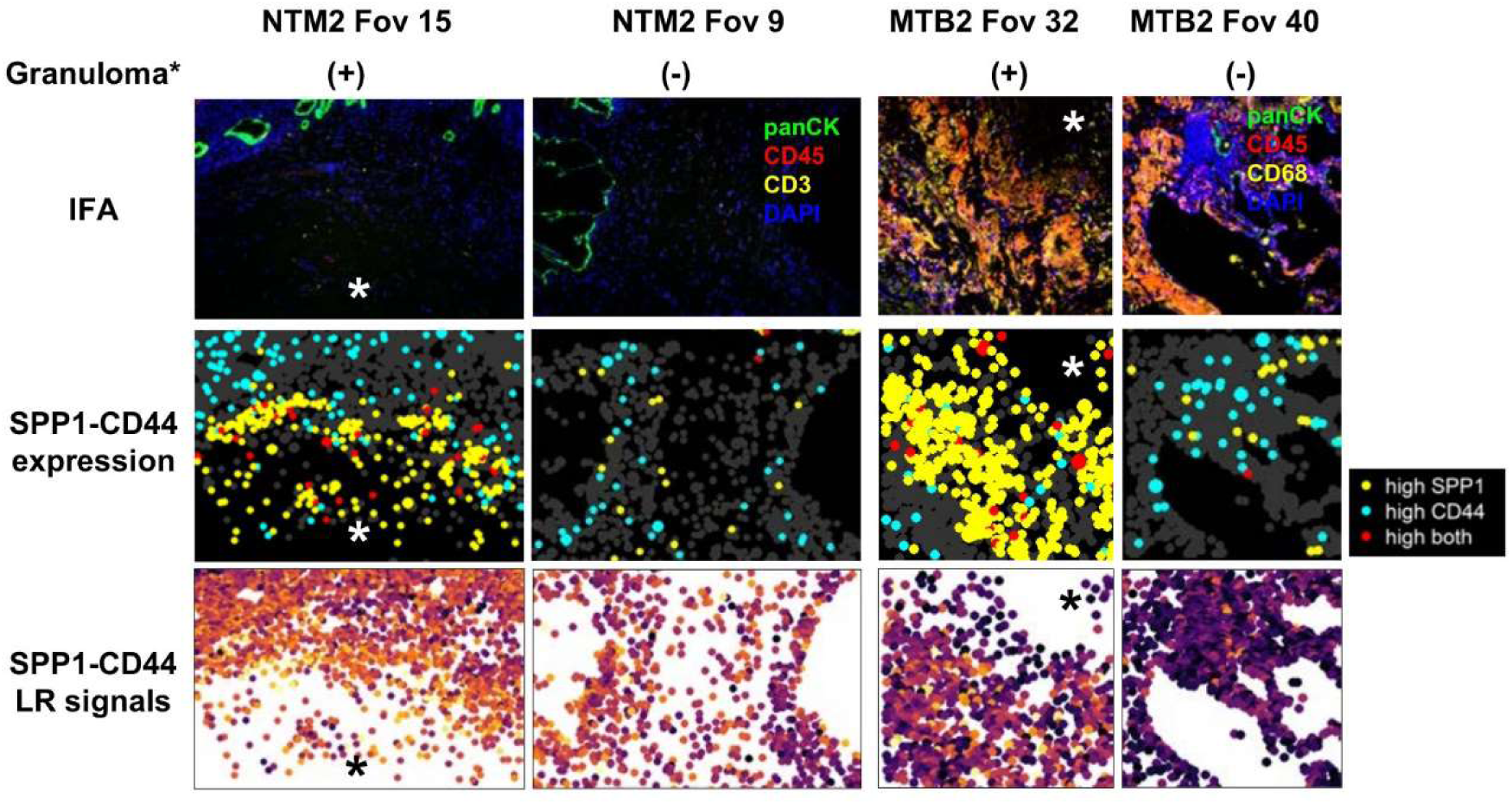
Co-expression and ligand-receptor assay in InSituCor. Immunofluorescence assay (IFA, upper panels) spatial images showing SPP1-CD44 expression (middle panels), and ligand-receptor (LR) signals of SPP1-CD44 (lower panels) in MTB and NTM tissues. In the SPP1-CD44 expression images, high expression of SPP1, CD44, and both are indicated by yellow, blue, and red dots, respectively. In the SPP1-CD44 LR signals images, yellow indicates high correlation. Spatial images from CosMx™ analysis were taken from fovs with (+) and without (−) granuloma in MTB or NTM samples. Asterisks (*) show the positions of granulomas.

**Supplementary Fig. 11:**
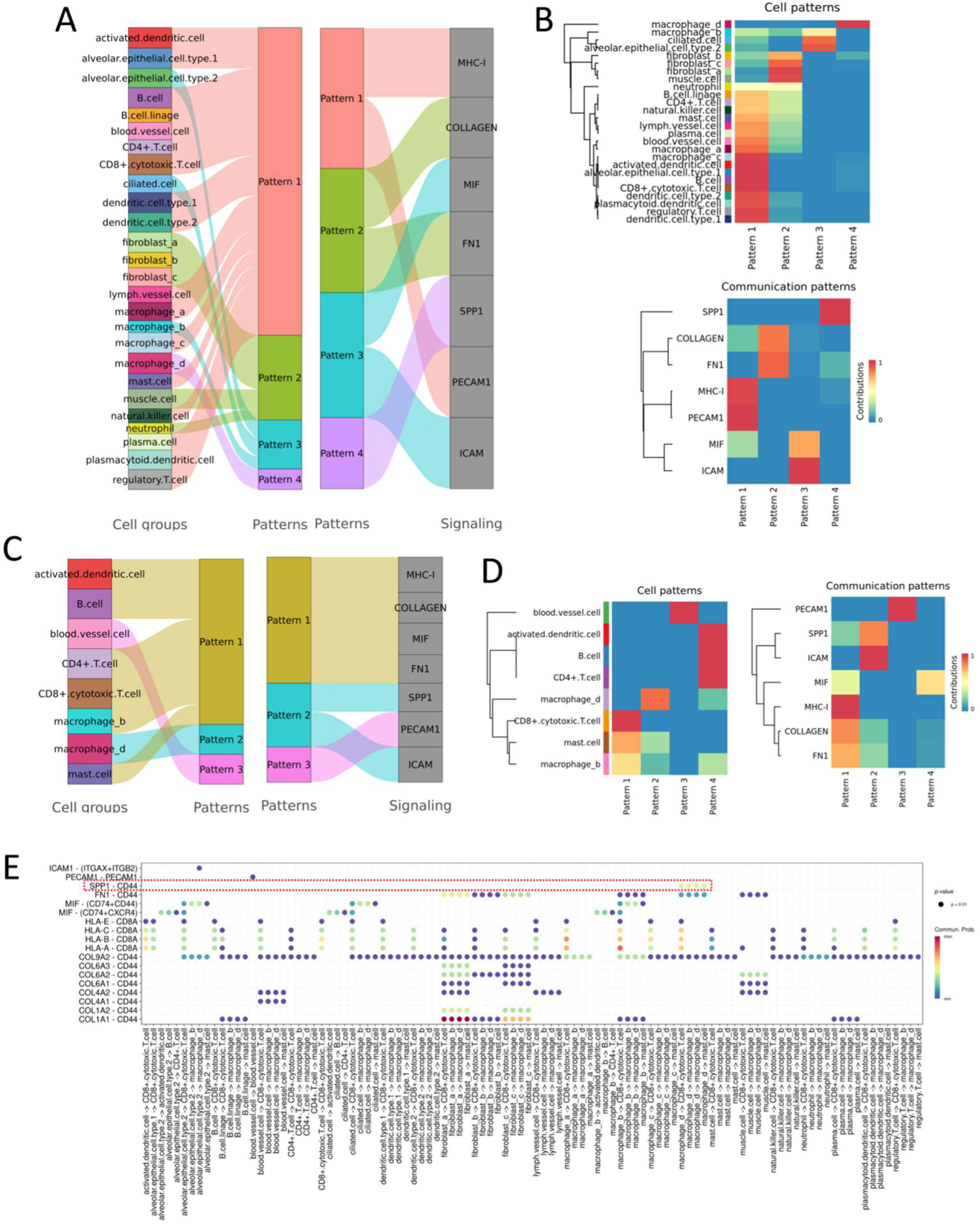
Outgoing and incoming communication patterns of secreting cells analyzed using CellChat in MTB and NTM samples. A. River plot showing cell patterns (left panel) and cell-communication patterns (right panel) for outgoing signaling pathways. B. Heatmaps displaying cell patterns (upper panel) and cell-communication patterns (lower panel) for outgoing signaling pathways. C. River plot showing cell patterns (left panel) and cell-communication patterns (right panel) for incoming signaling pathways. D. Heatmaps displaying cell patterns (upper panel) and cell-communication patterns (lower panel) for incoming signaling pathways. E. Dot plots illustrating the incoming and outgoing interaction strength of each cell-cell communication across all ligand-receptor interactions. The SPP1-CD44 signaling pathway is highlighted with a red dashed box.

**Supplementary Table S1:**
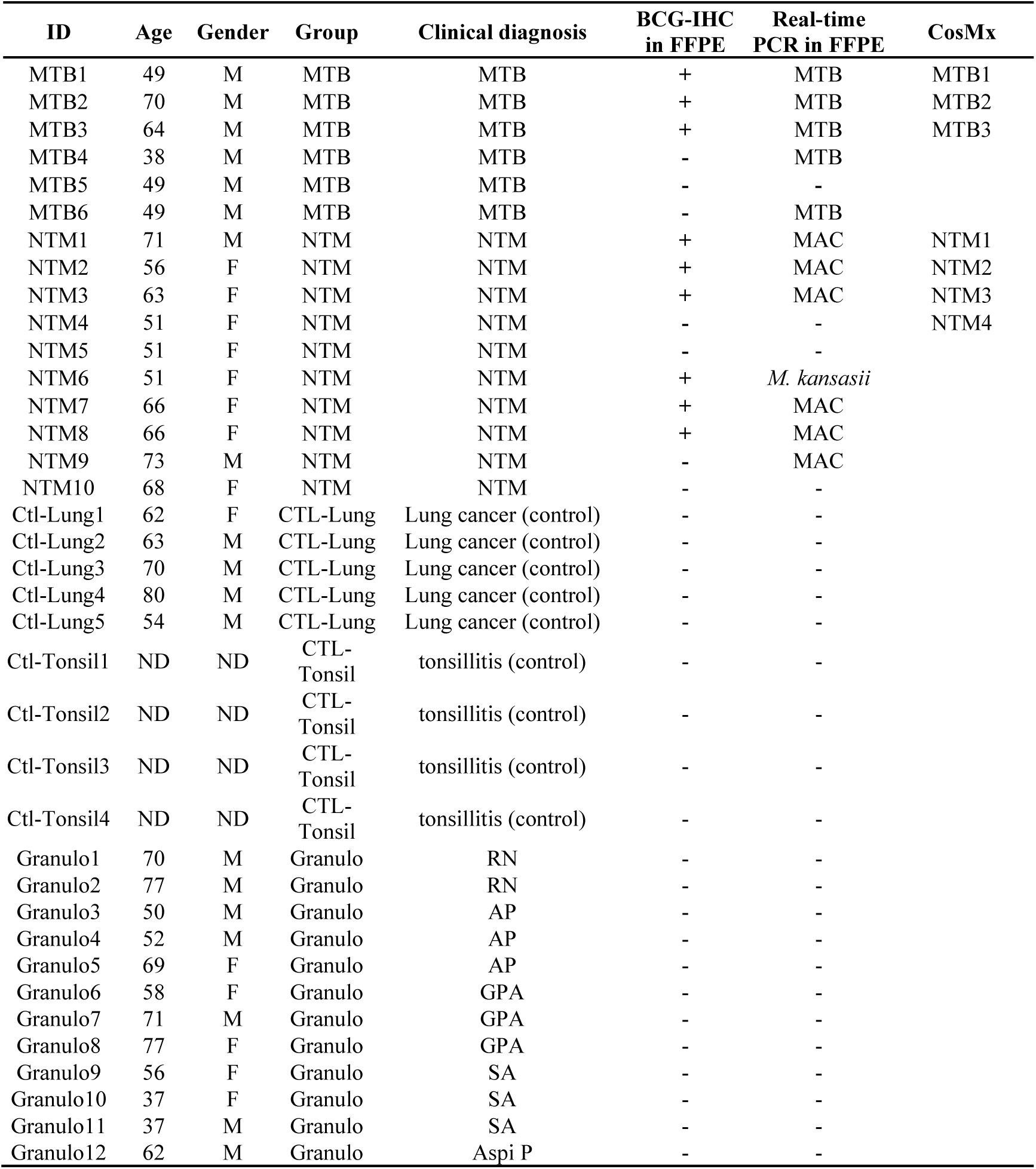
patient and sample list. AP: Aspergillus pneumonia, Aspi P: aspiration pneumonia, BCG: Bacille Calmette-Guerin, CTL: control, FFPE: formalin-fixed paraffin-embedded sample, GPA: granulomatosis with polyangiitis, IHC: immunohistochemistry, MAC: mycobacterium avium intracellulae complex, MTB: mycobacterium tuberculosis, NTM: non tuberculosis mycobacterium infection, ND: no data, RN: rheumatoid nodule, SA: sarcoidosis.

**Supplementary Table S2:**
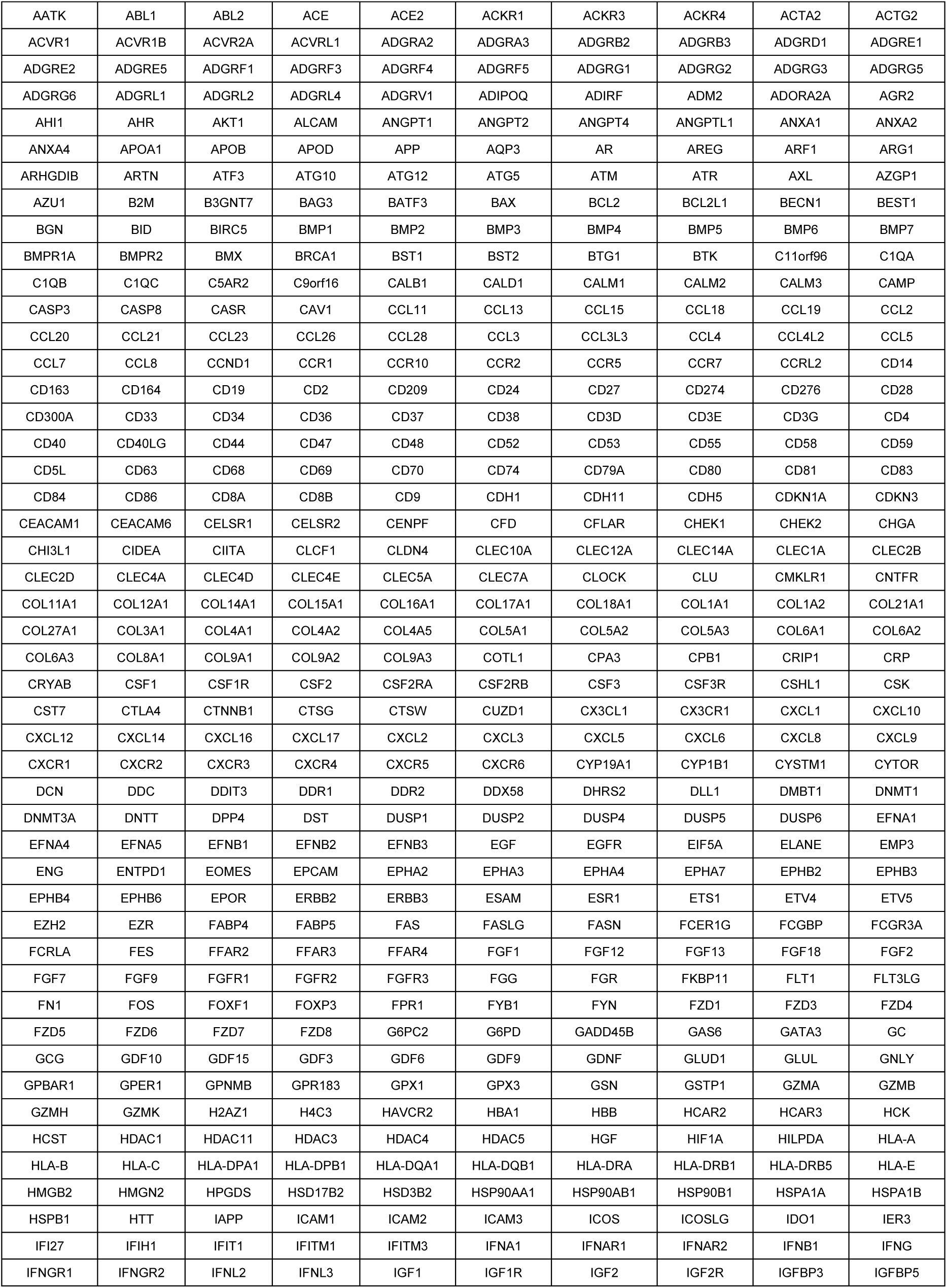
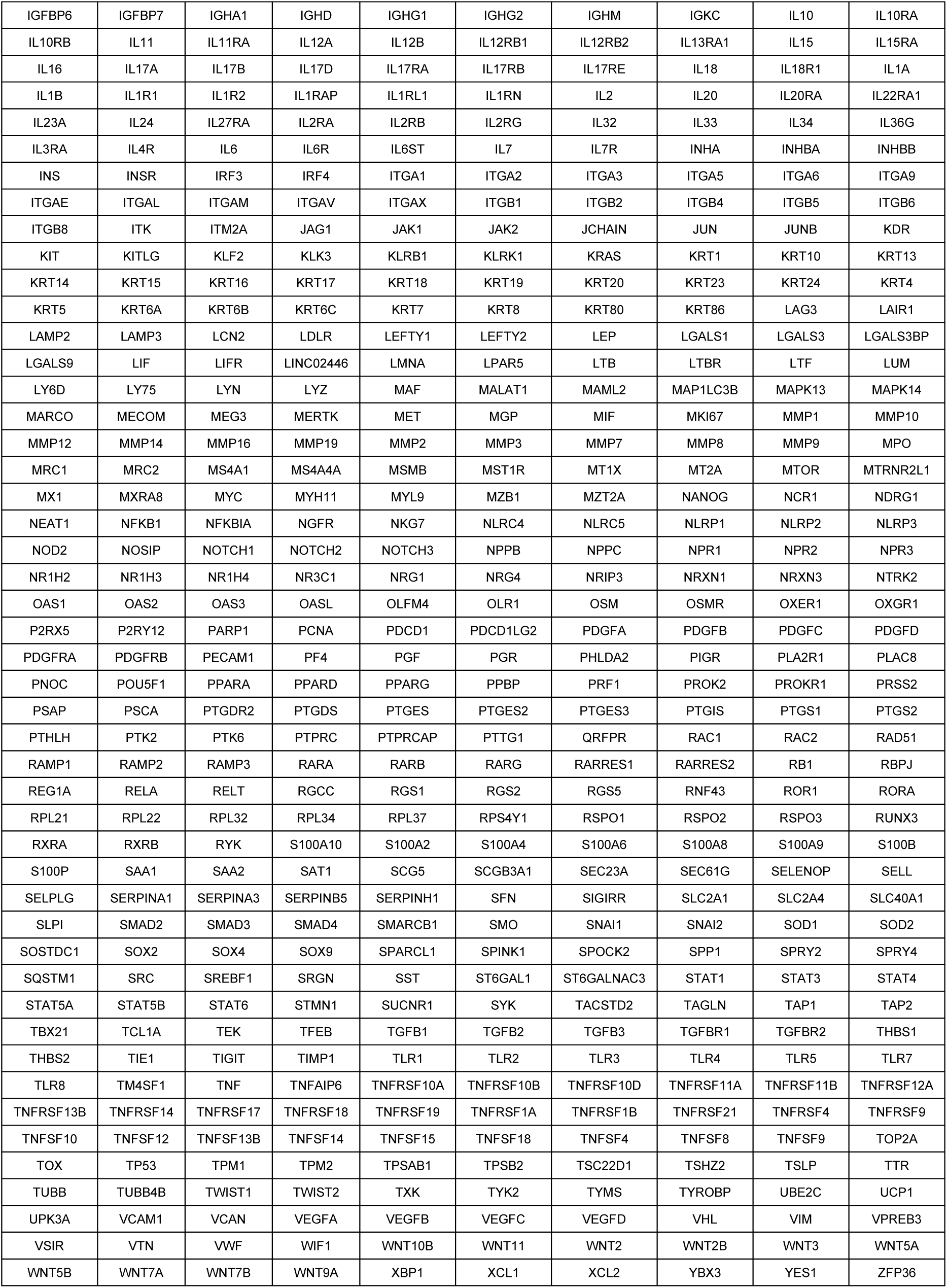
Gene list of the ‘CosMx Human Universal Cell Characterization’ panel.

**Supplementary Table S3:** Summary of CosMx assay on 7 slides of MTB and NTM samples. FOV: focus of view.

|  | Tissue | MTB1 | MTB2 | MTB3 | NTM1 | NTM2 | NTM3 | NTM4 | Total | Mean |
| --- | --- | --- | --- | --- | --- | --- | --- | --- | --- | --- |
| Number_of_FOVs |  | 21 | 47 | 45 | 21 | 15 | 22 | 45 | 216 | 30.9 |
| Total_Tissue_Area_mm2 |  | 8.33 | 4.65 | 4.16 | 3.63 | 2.09 | 4.28 | 5.08 | 32.2 | 4.6 |
| Number_of_Cells_in_Final_Analysis |  | 70649 | 40962 | 34799 | 31874 | 23692 | 51176 | 47689 | 300841 | 42977 |
| Total_Transcripts_Assigned_to_Cells |  | 8768373 | 7013069 | 4749647 | 4267383 | 1882347 | 5634413 | 7876583 | 40191815 | 5741688 |
| Mean_Transcripts_per_Cell |  | 124.1 | 171.2 | 136.5 | 133.9 | 79.5 | 110.1 | 165.2 | 920.5 | 131.5 |
| Maximum_Transcripts_per_Cell |  | 2226 | 2079 | 1697 | 2339 | 728 | 1362 | 2944 | 13375.0 | 1910.7 |
| Mean_Unique_Genes_Per_Cell |  | 70.14 | 65.82 | 58.73 | 81.5 | 54.69 | 69.42 | 79.18 | 479.5 | 68.5 |
| Mean_Transcripts_per_um2 |  | 0.0357 | 0.0222 | 0.0178 | 0.0458 | 0.0323 | 0.0466 | 0.0228 | 0.2 | 0.0 |

**Supplementary Table S4:** Cell population in CosMx assay on 7 slides of MTB and NTM samples.

| Cell types | MTB1 | MTB2 | MTB3 | NTM1 | NTM2 | NTM3 | NTM4 | overall_<br>percentage |
| --- | --- | --- | --- | --- | --- | --- | --- | --- |
| activated.dendritic.cell | 0.71 | 0.15 | 0.25 | 0.36 | 1.75 | 0.34 | 1.46 | 0.72 |
| alveolar.epithelial.cell.type.1 | 0.94 | 0.53 | 1.36 | 1.4 | 5.18 | 1.05 | 14.12 | 2.76 |
| alveolar.epithelial.cell.type.2 | 6.48 | 8.67 | 4.26 | 9.7 | 5.88 | 7.49 | 3.95 | 6.72 |
| B.cell | 3.97 | 10.71 | 6.36 | 19.78 | 9.82 | 2.82 | 6.77 | 7.87 |
| B.cell.lineage | 5.63 | 17.05 | 16.73 | 5.54 | 11.32 | 14.11 | 10.89 | 11.19 |
| blood.vessel.cell | 3.38 | 2 | 1.62 | 6.5 | 6.13 | 4.81 | 5.28 | 4.16 |
| CD4+.T.cell | 3.98 | 5.24 | 5.39 | 6.38 | 1.4 | 5.09 | 0.97 | 4.07 |
| CD8+.cytotoxic.T.cell | 3.24 | 1.42 | 1.76 | 2.75 | 7.23 | 1.2 | 3.16 | 3.12 |
| ciliated.cell | 0.88 | 0.1 | 0.24 | 0.84 | 0.39 | 0.22 | 4.96 | 0.83 |
| dendritic.cell.type.1 | 1.06 | 2.68 | 1.86 | 1.49 | 1.1 | 0.69 | 2.05 | 1.45 |
| dendritic.cell.type.2 | 1.71 | 3.13 | 4.55 | 5.09 | 1.64 | 4.15 | 1.5 | 2.95 |
| fibroblast_a | 1.07 | 4.53 | 4.58 | 0.61 | 4.46 | 4.59 | 5.24 | 3.36 |
| fibroblast_b | 2.06 | 11.41 | 17.16 | 17.7 | 16.12 | 19.73 | 10.74 | 12.61 |
| fibroblast_c | 21.16 | 1.29 | 1.69 | 0.37 | 0.43 | 2.82 | 0.84 | 5.97 |
| lymph.vessel.cell | 1.48 | 1.81 | 1.21 | 0.95 | 0.89 | 0.78 | 0.81 | 1.18 |
| macrophage_a | 15.32 | 0.16 | 0.24 | 0.14 | 0.2 | 1.17 | 0.84 | 3.95 |
| macrophage_b | 12.67 | 0.11 | 0.3 | 0.53 | 0.63 | 0.89 | 0.35 | 3.36 |
| macrophage_c | 0.86 | 6.17 | 3.45 | 1.53 | 7.26 | 5.11 | 9.36 | 4.39 |
| macrophage_d | 0.25 | 2.36 | 6.79 | 7.16 | 5.03 | 1.16 | 2.78 | 3.18 |
| mast.cell | 0.95 | 0.42 | 0.28 | 1.33 | 0.97 | 0.95 | 0.92 | 0.84 |
| muscle.cell | 0.31 | 1.14 | 0.64 | 2.04 | 2.67 | 0.45 | 2.92 | 1.27 |
| natural.killer.cell | 0.99 | 3.42 | 6.39 | 1.16 | 1.91 | 7.19 | 3.5 | 3.3 |
| neutrophil | 1.17 | 1.39 | 2.53 | 2.44 | 0.38 | 6.36 | 1.11 | 2.18 |
| plasma.cell | 1.51 | 11.47 | 8.01 | 1.36 | 3.1 | 4.33 | 2.21 | 4.38 |
| plasmacytoid.dendritic.cell | 1.61 | 2.38 | 1.94 | 2.08 | 1.59 | 2.08 | 1.3 | 1.85 |
| regulatory.T.cell | 6.61 | 0.24 | 0.4 | 0.75 | 2.51 | 0.41 | 1.96 | 2.36 |

